# IMK2-IMK3 module monitors biogenesis of nascent cell walls in land plants

**DOI:** 10.1101/2025.07.09.662438

**Authors:** Tetyana Smertenko, Deirdre Fahy, Glenn Turner, Andrei Smertenko

**Affiliations:** Institute of Biological Chemistry, Washington State University, Pullman, WA 99164, USA

## Abstract

Cell walls in land plants originate during cytokinesis from a membrane compartment called the cell plate. Arising as the trans-Golgi compartment, each cell plate acquires a unique molecular identity of lipids and proteins, associates with microtubules, and accumulates oligosaccharides in its lumen. Oligosaccharide deposition triggers microtubule depolymerisation and initiates a transition toward the molecular identity of the plasma membrane. How cells monitor cell plate biogenesis and coordinate it with microtubule behavior remains unknown. Here, we identify a novel signalling module composed of the receptor-like kinases IMK2 and IMK3, which integrates TRAPPII- and exo-cyst-dependent vesicle trafficking, the anti-parallel architecture and stabilization of microtubules, and the synthesis of oligosaccharide callose. Our findings reveal a mechanism for monitoring biogenesis of nascent cell walls.

## Introduction

Plant cell walls provide mechanical support to tissues and organs, facilitate the transport of water and solutes throughout the plant body, and protect against both abiotic and biotic stresses. Cell wall development occurs in three stages: (i) nascent cell wall formation; (ii) primary cell wall synthesis; and (iii) cell wall differentiation. Nascent walls are formed during cytokinesis as the cell plate – a two-dimensional array of cytoplasmic membrane compartments containing oligosaccharides in their lumen (Bajer, 1968, Hepler & Jackson, 1968). Subsequent chemical and structural modifications of the cell plate during tissue differentiation give rise to the full functional diversity of cell walls. Cell plates are produced by a plant-specific secretory module known as the phragmoplast, which comprises cytoskeletal elements, membrane compartments, and associated proteins (Smertenko *et al*., 2017, Bajer, 1968, Hepler & Jackson, 1968).

Cell plate biogenesis encompasses a series of evolutionarily conserved morphological and chemical transitions (Segui-Simarro *et al*., 2007). The phragmoplast guides trans-Golgi-derived vesicles carrying cell plate building blocks along microtubule scaffolds to the cell plate assembly location in the midzone (**Figure 1A**,**B**). There, vesicles fuse and remodel to form dumbbell-shaped compartments, which progressively fuse and reshape into a tubule-vesicular network. At this stage, the cell plate acquires a unique composition of lipids and membrane proteins, establishes connections with microtubules, and accumulates the oligosaccharide callose in its lumen. Callose accumulation triggers microtubule depolymerization and initiates the acquisition of plasma membrane identity, followed by the replacement of callose with common primary cell wall oligosaccharides: cellulose, hemicellulose, and pectin (Muller & Jurgens, 2016). Thus, the timely transition in cell plate identity requires coordination between chemical changes in the lumen and cytoplasmic processes, most likely mediated by a sensor with a receptor domain facing the lumen and a signalling domain facing the cytoplasm (**Figure 1B**). Here, we identify the molecular mechanism that ensures fidelity in the sequential stages of cell plate biogenesis, culminating in its hallmark identity transition – microtubule depolymerization.

**Fig. 1.**
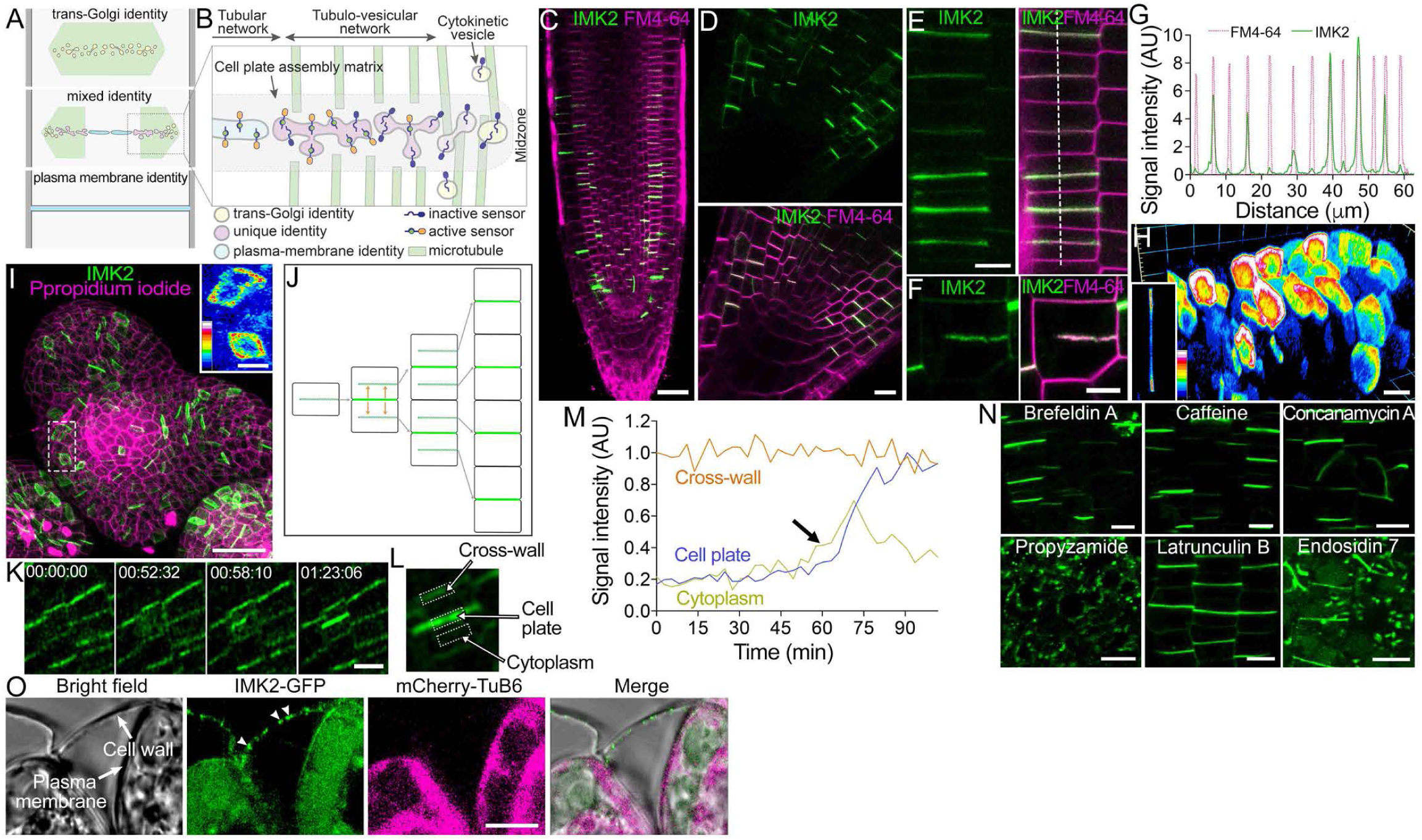
Characterization of Arabidopsis IMK2. A, Identities of nascent cell wall compartments. **B,** A sensor promotes microtubule depolymerization in response to oligosaccharide deposition. **C,D,** IMK2-GFP in the root apical meristem **(C;** scale bar, 30 µm) and root tip **(D;** scale bar, 10 µm). **E,F,** IMK2-GFP accumulates at alternate cross walls **(E)** and the cell plate **(F)** in the root apical meristem. Scale bars, 10µm. **G,** Normalized fluorescence intensities of IMK2-GFP and FM4-64 measured along the line in **E.** H, 3D-reconstruction showing enrichment of IMK2-GFP at cross wall edges. Scale bar, 10 µm. **I,** Maximum projection of IMK2-GFP in the inflorescence meristem. Scale bar, 20 µm. Inset shows the magnified region. Scale bar, S µm. **J,** A model illustrating IMK2 relocation from the parental cross wall to the cell plate. **K,** Time-lapse series showing dynamics IMK2-GFP during cell division. Numbers indicate relative time in hours:minutes:seconds. Scale bar, 20 µm. **L,M,** IMK2-GFP fluorescence intensity in three regions **(L)** normalized to the cross wall signal in frame 0 **(M)**. N, Localization of IMK2-GFP in the root apical meristem after three-hour treatment with 10 µM brefeldin A, 1 µM concanamycin A, 100 µM caffeine, S µM latrunculin B, 20 µM propyzamide, or 40 µM endosidin 7. Scale bars, 10 µm. **O,** IMK2-GFP associates with the cell wall (arrowheads) in plasmolized BY-2 cells. Scale bar, 10 µm.

### Receptor like kinase IMK2 governs cell division

Plants sense chemical changes in the extracellular space using receptor-like kinases (Kinoshita *et al*., 2005, ten Hove *et al*., 2011). A gene encoding a member of the *Arabidopsis* leucine-<u>r</u>ich <u>r</u>epeat domain <u>r</u>eceptor-like <u>k</u>inase family (LRR-RLK), the Inflorescence <u>M</u>eristem receptor-like <u>K</u>inase 2 (IMK2), co-expresses with several known regulators of cell division (**Figure S1A**). IMK2 domain architecture includes a unique N-terminal signal peptide, and conserved extracellular leucine rich repeat domain, a trans-membrane domain, and a cytoplasmic kinase domain (**Figure S1B**).

We isolated two homozygous T-DNA knockout alleles of *imk2* in the Col-0 background: *imk2-1* and *imk2-2* (**Figure S2A-C**)(ten Hove et al., 2011). Although the plants showed no discernible developmental or root growth aberrations, the root apical meristems of both alleles were shorter relative to the controls (**Figure S2D-H**). Expression of IMK2-GFP under the control of the endogenous promoter (*proIMK2:IMK2-GFP*) in *imk2-2* rescued the short meristem length phenotype in three independent transgenic lines (**Figure S2I**). GFP fluorescence of *imk2-2*;*proIMK2:IMK2-GFP* lines was stronger in the root apical meristem than in the elongation zone (**Figure S3A,C**). Another member of the LRR-RLK family BRI1-GFP expressed under its endogenous promoter and used as a control, showed strong signal in both the root apical meristem and the elongation zones (**Figure S3B,C**).

IMK2-GFP was enriched in the cell plates and newly formed cell walls in all tissues of the root apical meristem, except for the quiescent centre (**Figure 1C**,**D**). Every second cross wall appeared brighter (**Figure 1E**-**G**) whereas BRI1-GFP localized evenly on all cross walls and lateral walls (**Figure S3D**). IMK2-GFP accumulated at the cross wall attachment site to the lateral wall (**Figure 1H**). Expression of IMK2 under the control of the constitutive CaMV35S promoter resulted in its localization to both the lateral and cross walls (**Figure S3E**) indicating that *IMK2* expression level is critical for its polar localization.

We compared localization of IMK2 with two other key cytokinetic proteins: the syntaxin KNOLLE, responsible for membrane delivery to the cell plate, and the dynamin-related protein DRP1a, which contributes to cell plate remodelling (Lukowitz *et al*., 1996, Fujimoto *et al*., 2008, Kang *et al*., 2003). DRP1a localized to the cross wall only during cytokinesis and KNOLLE labelled only some cross walls (**Figure S3F-I**). The distinct localizations suggest that IMK2 is unlikely to function in vesicle delivery or membrane remodelling.

IMK2-GFP labelled cross walls in all growing and differentiating organs examined: inflorescence meristem (**Figure 1I**), cotyledonary and heart-shaped embryos, developing style, stomata, and lateral root primordia (**Figure S3J-O**). No signal was detected in differentiated tissues. In all cases IMK2-GFP accumulated at the cell plate edges, as illustrated by the inflorescence meristem (inset in **Figure 1I**).

The accumulation on IMK2-GFP at every second cross wall could be driven by IMK2 recycling from the parental cross wall to the cell plate during cytokinesis (**Figure 1J**). To test this hypothesis, we measured the IMK2-GFP signal at the cell plate, the cross wall, and the cytoplasm during cell division (**Figure 1K,L; Movie S1**). IMK2-GFP signal at the cross-wall remained steady throughout the cell division, while the cytoplasmic signal increased prior to the cell plate formation marked by the arrow in **Figure 1M**. Accumulation of IMK2-GFP at the cell plate coincided with reduction of the cytoplasmic signal. Thus, the bulk of IMK2 at the cell plate is synthesized de novo.

Next, we identified factors that maintain IMK2 localization at the cross walls. To this end, we applied inhibitors targeting microtubule polymerization (propyzamide), actin polymerization (latrunculin B), vesicle trafficking (brefeldin A, concanamycin A, and caffeine), and the synthesis of oligosaccharide callose (endosidin 7). Only propyzamide and endosidin 7 caused displacement of IMK2 from the cross walls and its accumulation in cytoplasmic bodies (**Figure 1N**).

The loss of cross wall localization upon inhibition of callose synthase suggests that IMK2 binds to the cell wall. Testing the association of proteins with cell walls in organs is challenging. Therefore, we generated a single-cell model by expressing *A. thaliana* IMK2-GFP and a microtubule marker mCherry-TuB6, both under the control of the CaMV35S promoter in tobacco BY-2 tissue culture cells. IMK2-GFP localized to the cell plate and accumulated at the attachment sites to the parental cell wall (**Figure S4A**-**C; Movie S2**). Cytokinetic vesicles delivering IMK2 were visible in the phragmoplast leading zone (**Figure S4D**). Although inhibition of callose synthesis by endosidin 7 allowed cell plate assembly as revealed by staining with FM4-64, the IMK2 signal was missing at the newly formed cell plate regions (**Figure S4E**). As shown in **Figure S4F** and **Movie S3**, IMK2-GFP is delivered to the cell plate but only a very weak GFP signal remains behind the expanding phragmoplast. The localization of another cell plate protein, the formin homology domain 8 (FH8), was not affected (**Figure S4E**). Hence, IMK2 localization in the cell plate in BY-2 cells requires callose synthesis.

To examine binding of IMK2 to the cell wall, we induced plasmolysis by supplementing tissue culture medium with 0.6 M mannitol. The osmotic shock caused depolymerization of microtubules resulting in a cytoplasmic tubulin signal (**Figure 1O**). However, IMK2-GFP remained on the cell walls, as well as on the plasma membrane, and in the cytoplasm.

IMK2 binds more strongly to the cell plate during the early biogenesis stages, consistent with its role in sensing cell plate assembly. Fluorescence loss in photobleaching (FLIP) showed no reduction of the fluorescence signal at the cell plate edges (region E) whereas signal in the centre (region C) declined (**Figure 2A,B)**. We measured the ratio of GFP signal intensity between regions of interest E at the cell plate edge and C at the centre (I^E^/I^C^; **Figure 2A**) after 120 seconds of the experiment. While the IMK2-GFP signal decayed slower in the cell plate edges relative to the centre, FH8-GFP showed similar signal decay in both regions (**Figure 2B**,**C**). Persistence of the IMK2-GFP signal indicates a stronger association with the cell plate.

**Fig. 2.**
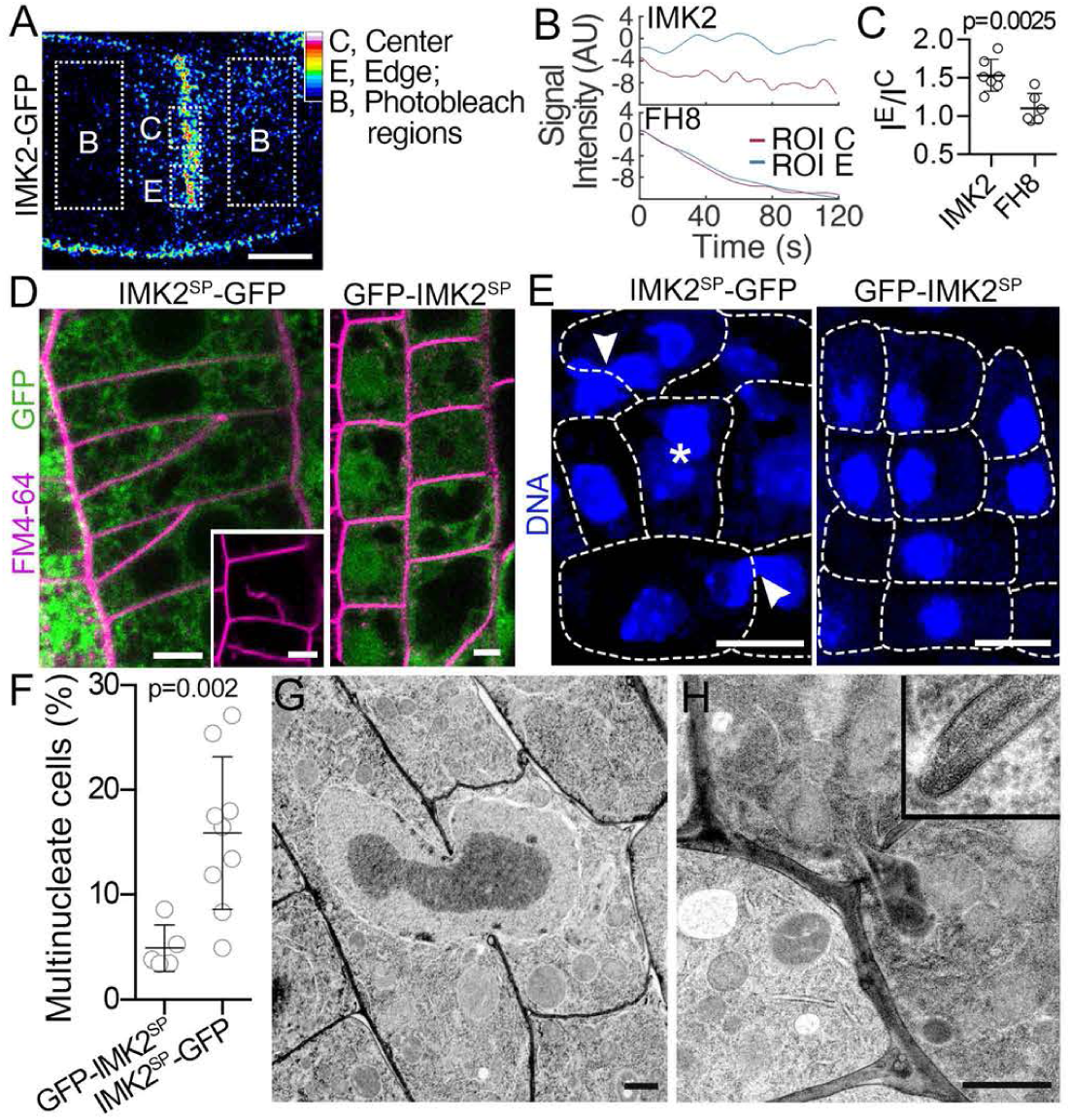
IMK2 is important for cell division. A, Schematic of the FLIP experiment using IMK2-GFP as an example. Photobleaching was applied in ROIs 8, and signal intensity was monitored in ROIs C and E. Scale bar, 10 µm. **B,C,** Signal decay in FLIP experiments for IMK2-GFP and FHS-GFP **(B),** and the ratio of signal intensity in ROIs E and C, JE and Jc, after 120 s (C). p-value calculated by an unpaired *t-test* (n=7). **D,** Expression of IMK2^SP^-GFP disrupts cell division, causing mispositioned cross walls and formation of cell wall stubs (inset) in root apical meristem cells. In contrast, cell division remains normal in cells expressing GFP-IMK2^SP^. Scale bars, 5 µm. **E,** DAPI staining reveals multinucleated cells (asterisk) and incompletely separated nuclei (arrowheads) in roots expressing IMK2^SP^-GFP, while no defects are observed in roots expressing GFP-IMK2^SP^. Scale bars, 10 µm. **F,** Frequency ofmultinucleated cells in the root apical meristems shown in **B.** p-value calculated by an unpaired *t-test* (n>S roots). **G,H,** Representative images of a nucleus **(G)** and a plastid **(H)** shared between incompletely divided cells in roots expressing IMK2^SP^-GFP. The inset shows intact plasms membrane at the cell wall fenestra. Scale bars, 1 µm.

A dominant-negative approach uncovered an essential role for IMK2 in cell division. Expression of the signal peptide with the first leucine-rich repeat fused to GFP (IMK2^SP^-GFP) under the control of CaMV35S promoter failed to produce viable transgenic plants, indicating that the construct was lethal. Expressing IMK2^SP^-GFP under the control of an estrogen-inducible promoter helped to overcome this bottleneck. Induction of IMK2^SP^-GFP expression for 72 hours resulted in cytokinetic failure (formation of stubs) and misplaced cross walls (**Figure 2D** left). DNA staining in root apical meristems with DAPI revealed a high frequency of multi-nucleate cells (**Figure 2E** left and **F**). Some cells shared an incompletely separated nucleus (**Figure 2E** left). The negative control plants expressing the N-terminal GFP-fusion of the signal peptide (GFP:IMK2^SP^) showed no cytokinetic defects (**Figure 2D** right and **E** right). Electron microscopy of IMK2^SP^-GFP root meristems confirmed the presence of shared nuclei and other organelles (e.g. plastids) between daughter cells (**Figure 2G**,**H**). Cytokinetic failure in IMK2^SP^-GFP lines, accumulation of IMK2-GFP at the cross walls, dependency on callose, association with the cell wall, and retention at the cell plate edges, demonstrate that IMK2 functions as a sensor of cell plate biogenesis.

### IMK2 forms a complex with IMK3

The mild phenotype of *imk2* knockout alleles suggests functional redundancy of IMK2. Receptor kinase signalling requires interactions with other receptor kinases, and IMK2-interacting partners could be responsible for the redundancy. To test this hypothesis, we isolated *A. thaliana* IMK2-GFP-interacting proteins by immunoprecipitation with anti-GFP from the BY-2 cell cultures synchronized to contain 42% of telophase cells. Considering that frequency of cytokinetic cells in root tips is between 0.1% and 0.01%, and tobacco cell plates are on average six-fold larger, the sensitivity of our approach for capturing IMK2 partners in cytokinesis is at least 2,500-fold greater than using *A. thaliana* lines. GFP antibodies detected a band of 120 kDa in the total protein extract and a 180 kDa band in the immuno-precipitated material (**Figure S5A**). No signal was detected in the negative control containing beads without GFP antibody. Proteomics analysis of the immunoprecipitated material identified two peptides corresponding to tobacco IMK3 with 99% and 100% confidence (**Figure S6**). Two other peptides in the pull down material corresponded to tobacco IMK2.

Expression of *A. thaliana* IMK3-GFP in BY-2 cells resulted in cell plate labelling (**Figure S5B**). Immunoprecipitation of IMK3-GFP from the cell-cycle-synchronized cells (40% telophase) identified tobacco IMK2 and IMK3 peptides (**Figure S6**).

The immunoprecipitation outcomes were verified with Bimolecular Fluorescence Complementation (BiFC). Co-expression of IMK2 fused to the N-terminal fragment of YFP with IMK3 fused to C-terminal fragment of YFP resulted in reconstitution of fluorescence at the plasma membrane (**Figure S5C**). Furthermore, the fluorescence was reconstituted in cells co-expressing IMK2 fused to YFP N-terminus and IMK2 fused to YFP C-terminus or IMK3 fused to YFP N-terminus and IMK3 fused to YFP C-terminus (**Figure S5D, E**). The N-terminal fusions of the YFP failed to reconstitute the signal (**Figure S5F, G**). As the N-terminal YFP fragment would interfere with IMK2 or IMK3 trafficking, this result indicates that the complex forms only in the plasma membrane. To examine whether interaction between IMK2 and IMK3 is a consequence of random collisions at the plasma membrane, we co-expressed IMK2 or IMK3 with an integral membrane formin AtFH1 or AtFH5. Here, the fluorescence was not reconstituted (**Figure S5H-K**). These results are consistent with the model whereby IMK2 and IMK3 form homo- and hetero-oligomers in the plasma membrane.

To examine the functions of IMK3, we characterized the phenotype of a homozygous *IMK3* T-DNA insertion allele in the Col-0 background, designated *imk3-1* (**Figure S7A**,**B**)(ten Hove et al., 2011). The root apical meristem of *imk3-1* was reduced in size compared to the Col-0 control; however mature plants appeared morphologically normal with comparable root growth to the Col-0 (**Figure S7C**-**F**). Transformation of *imk3-1* with *proIMK3:IMK3-GFP* rescued the meristem size phenotype in three independent transgenic lines (**Figure S7G**).

Analysis of *imk3-1;proIMK3:IMK3-GFP* plants shows that *IMK3* is expressed in the root apical meristem and accumulates in cell plates, newly formed cell walls (**Figure 3A**-**D**), and at cross wall attachment sites to the parental cell wall (**Figure 3E**). IMK3 localized to cross walls in all developing organs analysed – including the floral meristem (**Figure 3F**), heart and torpedo embryos, and in anther fibres and style (**Figure S8A**-**F**) - but was absent in differentiated tissues. Disruption of callose synthesis by endosidin 7 or microtubule polymerization by propyzamide did not affect IMK3-GFP localization (**Figure S8G**).

**Fig. 3.**
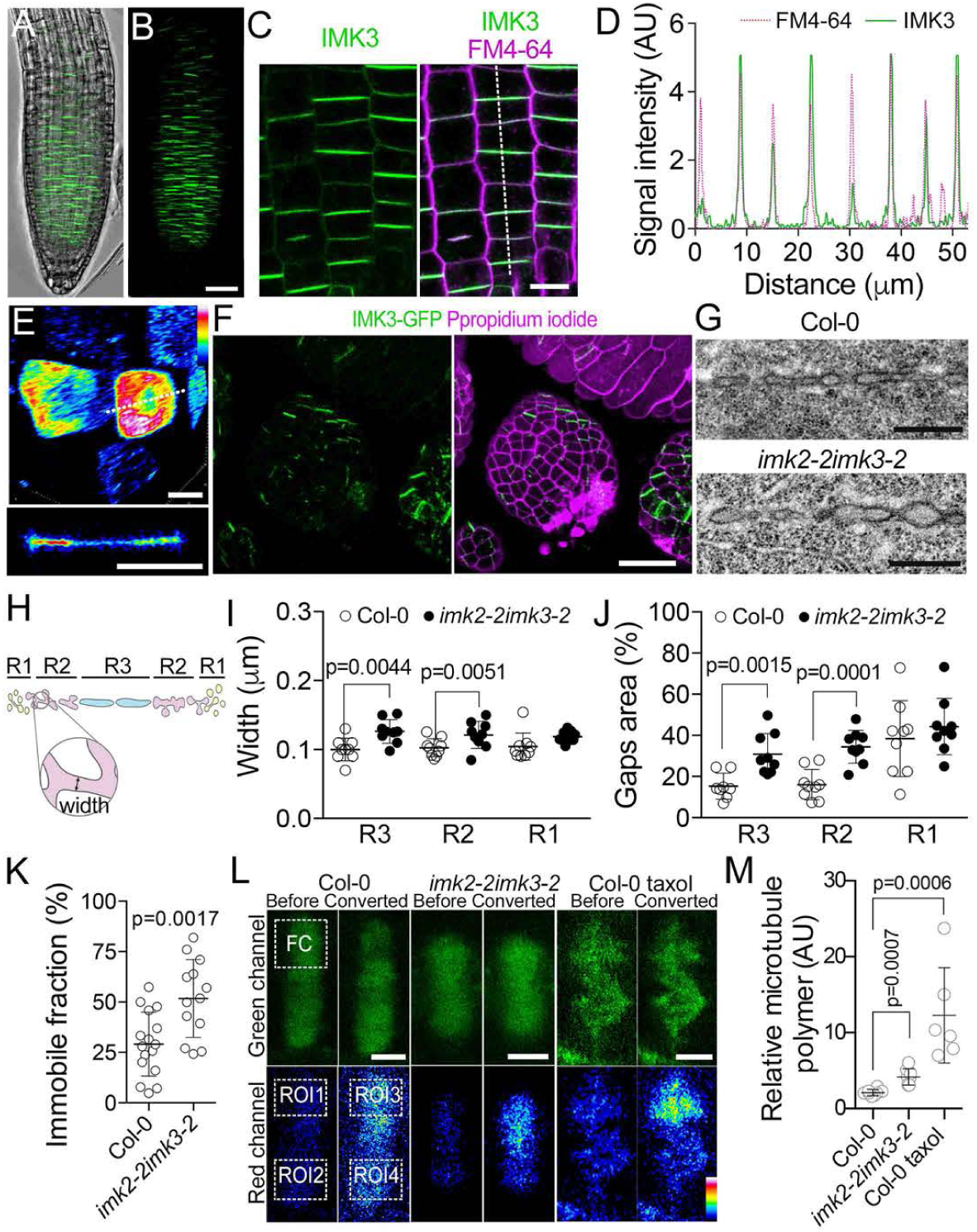
The IMK2-IMK3 module governs cytokinesis. A,B, Maximum projection of the root apical meristem in the *imk3-1;pro/MK3:IMK3-GFP* line. Scale bar, 40 µm. **C,** IMK3-GFP accumulates at alternate cross walls in the root apical meristem. Scale bar, 10 µm. **D,** Normalized fluorescence intensities of IMK3-GFP and FM4-64 measured along the line in **C.** E, 3D-reconstruction showing enrichment of IMK3-GFP at cross wall edges. Inset shows a single optical section along the white line. Scale bars, 5 µm. **F,** Maximum projection of IMK3-GFP in the inflorescence meristem. Scale bar, 20 µm. **G,** Representative transmission electron micrographs of cell plates in the root apical meristem of Col-0 and *imk2-2imk3-2* plants. Scale bars, 500 nm. **H-J,** Measurements of cell plate width **(I)** and gaps area **(J)** in three regions **(H)** of Col-0 and *imk2-2imk3-2* phragmoplasts. Nine cell plates per each genotype were analysed, with 30-35 compartments measured per cell plate. p-values calculated using an unpaired *t-test* (n=9). **K,** Immobile fraction of the mNeonGreen-TuB2 signal. p-values calculated using an unpaired *t-test (n=13)*. **L,** Representative images from EosFP-TuB2 photoconversion experiments. FC indicates the photoconverted region; fluorescence intensity was measured in ROIs 1-4. Scale bars, 3 µm. **M,** Relative abundance oftubulin polymer. p-values calculated using an unpaired *t-test (n=5)*.

Functional redundancy between *IMK2* and *IMK3* was tested by performing reciprocal complementation: expressing *proIMK3:IMK3-GFP* in *imk2-2* and *proIMK2:IMK2-GFP* in *imk3-1*. Expression of IMK3-GFP rescued the *imk2-2* apical meristem phenotype, whereas IMK2-GFP failed to rescue the *imk3-1* phenotype (**Figure S7H**). IMK3-GFP localization in the *imk2-2* background was indistinguishable from that in the control (**Figure 3C**), but IMK2-GFP in the *imk3-1* background localized to both cross walls and cytoplasmic vesicles (**Figure S7I**). Hence, IMK3 contributes to retaining IMK2 in the cell plate and IMK2 is not required for IMK3 localization.

### IMK2-IMK3 module regulates microtubule stability

To examine the function of the IMK2-IMK3 complex, we generated double mutants. As both genes are located proximally on chromosome 3, CRISPR/Cas9-mediated gene editing was used to mutagenize *IMK2* in *imk3-1* background, generating the *imk2-3imk3-1* allele, and to mutagenize *IMK3* in *imk2-2* background, generating the *imk2-2imk3-2* allele. Each allele contained an adenine insertion resulting in premature termination of the corresponding reading frame (**Figure S9A)**. The mutant plants appeared morphologically normal (**Figure S9B-D**). Both *imk2-2imk3-2* and *imk2-3imk3-1* alleles showed shorter primary roots (**Figure S9E**,**F)** and shorter root apical meristem (**Figure S9G**,**H**) relative to single mutant or Col-0 wild-type plants. Considering similarity of both alleles, all subsequent analyses were performed with the *imk2-2imk3-2* allele.

Phenotyping cell division in *imk2-2imk3-2* root apical meristem using transmission electron microscopy revealed a greater width of the membrane compartments relative to Col-0 control (**Figure 3G**). Next, we measured cell plate morphology in regions corresponding to vesicle delivery, fusion, and formation of tubule-vesicular network (region 1, R1), transition towards tubular network (region 2, R2), and the central part of cell plate (region 3, R3) (**Figures 3H** and **S10A**). The measurements were performed on the cross sections taken through the centre of cytokinetic cells, as confirmed by the middle cross-section through both daughter nuclei. The cell plate compartments in regions 2 and 3 were wider in *imk2-2cimk3-2* than in Col-0, whereas no differences were detected in region 1 (**Figure 3I**).

The gaps between cell plate compartments were assessed by measuring the percentage of cell plate area (coloured blue in **Figure S10B**) occupied by the membrane compartments (coloured black). The membrane compartments occupied a greater cell plate area in *imk2-2imk3-2* relative to the control in regions 2 and 3, but not in region 1 (**Figure 3J**). This means initial stages of the cell plate synthesis are not affected in *imk2-2imk3-2*, and defects arise during transition from the cell plate to the plasma membrane identity.

Microtubules appeared irregular in ca. 20% of *imk2-2imk3-2* phragmoplasts (**Figure S10C**), and phragmoplast expansion rate was slower, though the phragmoplast length was not affected (**Figure S10D,E**). We measured microtubule dynamics in the phragmoplast using fluorescence recovery after photobleaching (FRAP)(**Figure S11A**,**B**). Although the microtubule turnover rate was twofold faster in the mutant than in the Col-0 (**Figure S11C**,**D**), the immobile fraction in the mutant was twofold greater (**Figure 3K**). A combination of faster turnover rate and higher immobile fraction in the mutant indicates a greater abundance of microtubules relative to free tubulin.

As FRAP provides indirect assessment of the immobile fraction, we measured the fraction of stable microtubules directly using the photo-convertible tubulin probe EosFP:TuB2 (Schmidt-Marcec *et al*., 2023). The photo-conversion within a region of interest switches fluorescence of both microtubules and free tubulin from green to red (**Figure 3L**; **Movie S4**). Free tubulin diffuses throughout the cytoplasm within a second, whereas stable microtubules persist in the photoconverted region. The fraction of stable microtubules can be determined by taking the ratio of red signal within the converted region to that in the non-converted region. To test the sensitivity of this approach to changes in microtubule stability, we treated Col-0 control seedlings with the inhibitor of microtubule depolymerization, taxol. As expected, the fraction of stable microtubules was two-fold greater in the taxol treated cells than in the control. The fraction of stable microtubules in *imk2-2imk3-2* was 50% greater than in the Col-0 control (**Figure 3L,M**). As a negative control, we photo-converted EosFP-TuB2 in the region outside the phragmoplast and found no differences in fluorescence between different parts of the phragmoplast in Col-0 and *imk2-2imk3-2* (**Figure S11E,F**; **Movie S5**). From these results, we conclude that the IMK2-IMK3 module promotes micro-tubule depolymerization.

### IMK2-IMK3 module senses cell plate assembly

According to the current model, cell plate assembly starts with the establishment of antiparallel microtubule overlap in the phragmoplast midzone by MAP65 proteins (Smertenko, 2018). MAP65 interacts with TRAPPII tethering complex to facilitate recruitment of cytokinetic vesicles (Rybak *et al*., 2014). We tested the role of antiparallel overlaps in localization of IMK2 and IMK3 using a mild double knockout allele *map65-1map65-2* (Sasabe *et al*., 2011). Although IMK2 localized to the cross walls and the cell plate, some signal was detected in the vesicles (**Figure 4A**), and IMK2-GFP enrichment at the cross wall relative to the lateral wall was significantly reduced (**Figure 4E**,**F**). In contrast, the localization and cross wall enrichment of IMK3-GFP was not affected in *map65-1map65-2* (**Figure 4A**,**G**). Therefore, IMK2 recruitment to the midzone requires antiparallel microtubule bundling.

**Fig. 4.**
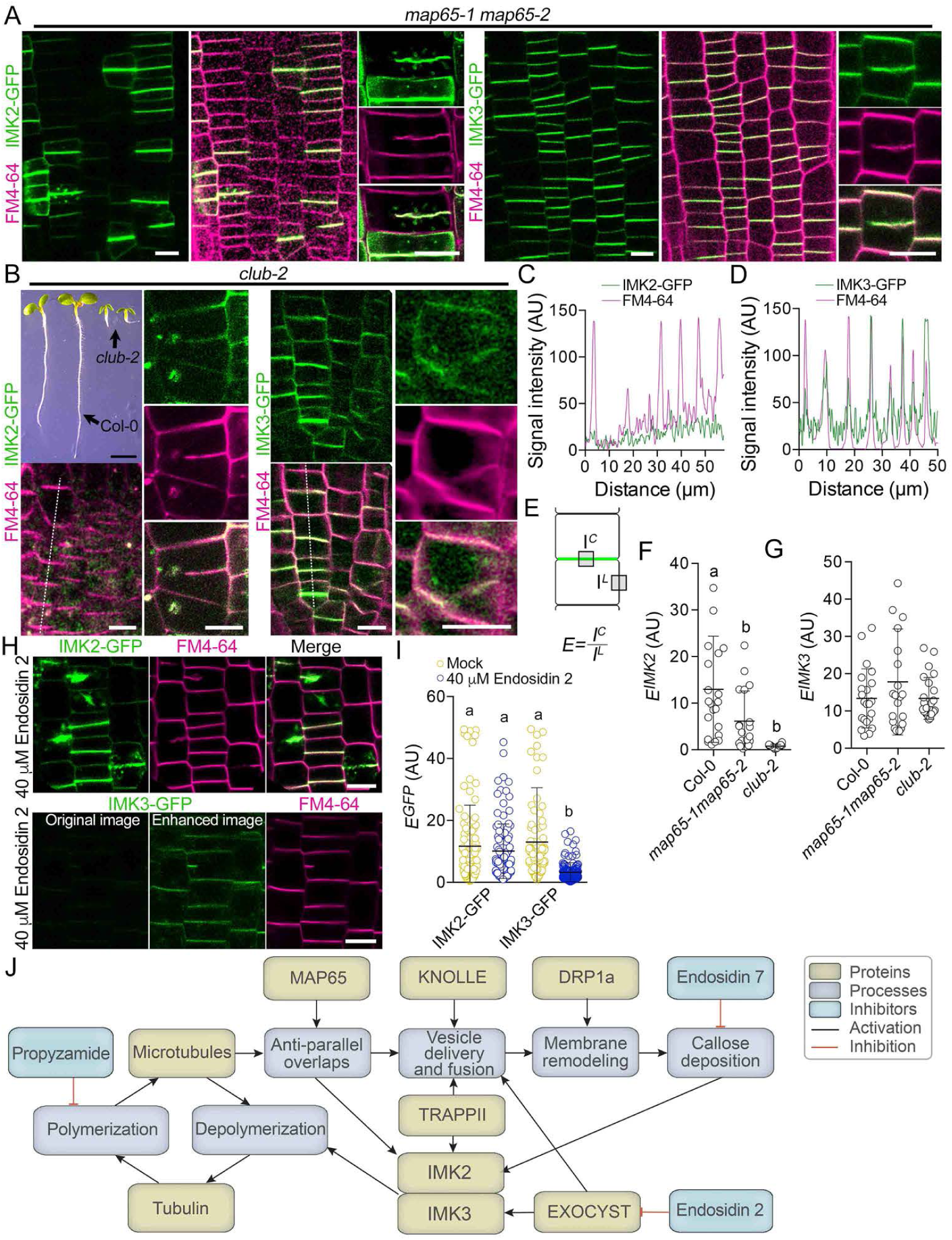
Different pathways deliver IMK2 and IMK3 to the cell plate. A, Localization of IMK2-GFP and IMK3-GFP in the *map65-1 map65-2* background. Scale bars, 10 µm. **B,** Localization of IMK2-GFP and IMK3-GFP in *club-2* background. Scale bar for the whole plant image, 5 mm, all others, 10 µm. **C,D,** Normalized fluorescence intensities of IMK2-GFP, IMK3-GFP, and FM4-64 measured along the indicated lines in **B.** E, Schematic illustration showing measurements of GFP signal enrichment Eat the cross walls. **F,G,** Quantification of IMK2-GFP **(F)** and IMK3-GFP **(G)** signal enrichment at cross walls in *map65-1map65-2* and *club-2* mutants. Datasets labelled with distinct letters are significantly different (p<0.05) according to one-way ANOVA **H,** Localization of IMK2-GFP or IMK3-GFP after three-hour treatment with 40 µm endosidin 2. Scale bars, 10 µm. **I,** Enrichment of IMK2-GFP and IMK3-GFP signals at cross walls in mock- or endosidin 2-treated cells. Datasets labelled with distinct letters are significantly different (p<0.05) according to one-way ANOVA **J,** Localization of IMK2-GFP at the cell plate depends on the formation of antiparallel overlaps in the midzone, TRAPPII-dependent vesicle trafficking, callose deposition, and the presence of IMK3. IMK3 localization is dependent on exocyst-mediated vesicle trafficking. Together, the IMK2-IMK3 complex promotes microtubule depolymerization.

As MAP65 functions upstream of TRAPPII-dependent vesicle tethering at the midzone, we analysed localization of IMK2 and IMK3 in the *club-2* allele of a gene encoding TRAPPII subunit TRS130 (Jaber *et al*., 2010). The homozygous *club-2* causes sterility, and the homozygous plants were identified in segregating population by the stunted growth phenotype (**Figure 4B**). IMK2 accumulation at the cell plate and the cross wall was lost in *club-2* (**Figure 4B,C**), and the signal distributed evenly on all cell walls and in vesicles. The enrichment of IMK2 at the cross walls was also lost (**Figure 4F**). IMK3 localized to the cell plates, cross walls, cell wall stubs, and lateral walls (**Figure 4B, D**), but the enrichment of IMK3 on the cross walls was not affected (**Figure 4G**).

It has been shown that TRAPPII functions together with the exocyst pathway during cytokinesis (Rybak et al., 2014). We examined the role of the exocyst in localization of IMK2 and IMK3 by treating roots with an inhibitor of plant exocyst complex, endosidin 2, for three hours. Consistent with the critical role of exocyst in cytokinesis, the treatment caused formation of cell wall stubs (**Figure 4H**). In the endosidin 2 treated cells, IMK2-GFP localized in cytokinetic vesicles, resembling that in *imk3-1* (**Figures 4H** and **S7H**). In contrast, the IMK3-GFP signal was very weak under the standard image acquisition conditions, but could be enhanced by image editing (**Figure 4H**). While the enrichment of IMK3-GFP on the cross walls was perturbed, the enrichment of IMK2-GFP was not affected (**Figure 4I**). These outcomes demonstrate that IMK2 and IMK3 exploit different trafficking pathways to reach the cell plate.

## Discussion

Here, we elucidate a quality control mechanism that governs cell plate biogenesis by monitoring three interconnected cellular processes: membrane trafficking, oligosaccharide (callose) synthesis, and cytoskeletal organization. Based on our findings and prior work, we propose the following model (**Figure 4J**). First, cell plate biogenesis begins with the formation of antiparallel microtubule overlaps in the phragmoplast midzone, mediated by MAP65 proteins (Smertenko et al., 2017, Smertenko, 2018). Second, cell plate building materials are delivered to the overlapping microtubules via TRAPPII- and exocyst-dependent vesicle tethering (Jaber et al., 2010, Rybak et al., 2014). Third, IMK2, along with other cargo, is delivered to the cell plate via TRAPPII-dependent vesicle tethering. Hence, IMK2 acts as a sensor of both antiparallel overlap fidelity and TRAPPII function, as disruptions in either process impair its delivery and retention. Fourth, IMK3 delivery to the cell plate is dependent on the exocyst complex. Fifth, both IMK3 and callose are required for IMK2 retention at the cell plate. Inhibition of callose synthesis or loss of *IMK3* disrupts IMK2 localization and function. Sixth, the IMK2-IMK3 complex promotes depolymerization of phragmoplast microtubules. Unless all processes responsible for targeting of the IMK2-IMK3 complex to the cell plate and callose synthesis function normally, microtubule depolymerization would fail leading to abnormal cell plate morphology, reduced cell plate synthesis rate, and impaired root growth.

The dependency of the IMK2-IMK3 module on multiple upstream processes positions it as a central hub in the network governing cell plate assembly fidelity. Given that phragmoplast microtubules in *imk2-2imk3-2* mutants can still depolymerize and the phragmoplast continues to expand, additional redundant pathways likely contribute to promoting microtubule destabilization. Characterizing these alternative pathways and elucidating their crosstalk with the IMK2-IMK3 module will be essential. IMK2 has been shown to interact with mitogen-activated protein kinases MAPK6 and MAPK8 (Popescu *et al*., 2009), and MAPK signalling is implicated in microtubule destabilization through phosphoregulation of microtubule-binding proteins MAP65 and End-Binding 1 (EB1) (Smertenko, 2025, Sasabe *et al*., 2006, Kohoutová *et al*., 2015). Several phosphorylation sites on MAP65 proteins remain unassigned to specific signalling pathways and may be regulated by the IMK2-IMK3 module (Smertenko, 2025). Other potential substrates include formins, which contribute to microtubule anchoring at the cell plate (Zhang *et al*., 2021). Intriguingly, IMK2 and IMK3 persist in the cross walls beyond cytokinesis. Considering that IMK2 interacts with several known and unknown receptor-like kinases (Smakowska-Luzan *et al*., 2018), the IMK2-IMK3 module may also contribute to cell wall differentiation.

## Materials and methods

### Constructs

All primers used in this study are provided in **Table S1,** all constructs and lines are listed in **Table S2**, and the gene accession numbers are listed in **Table S5**. To express *IMK2* (At3G51740) or *IMK3* (At3G56100) under the CaMV35S promoter, full-length cDNAs were amplified using primers IMK2-GW-F and IMK2-GW-R or IMK3-GW-F and IMK3-GW-R, respectively. These fragments were cloned into Gateway binary vector *pMDC83* (Curtis & Grossniklaus, 2003) to generate C-terminal GFP fusion proteins. For *proIMK2:IMK2-GFP* and *proIMK3:IMK3-GFP* fusions, the genomic regions of *IMK2* (1,393 nt upstream of the ATG to the final codon) and *IMK3* (1,446 nt upstream of the ATG to the last codon) were amplified by PCR using primers pIMK2-IMK2-GW-F and IMK2-GW-R or pIMK3-IMK3-GW-F and IMK3-GW-R, respectively. The PCR fragments were first cloned into *pDONR207* and then into *pMDC107* (Curtis & Grossniklaus, 2003) using Gateway technology (Invitrogen). The proBRI1:BRI1-GFP was previously described (Geldner *et al*., 2007).

CRISPR/Cas9-mediated knockouts were generated by selecting target regions with no predicted off-targets and designing protospacers for cloning into the entry cassette, as described (Schiml *et al*., 2016). Complementary protospacer oligonucleotides for each of *IMK2* and *IMK3* (IMK2-Sa-sgRNA-F and IMK2-Sa-sgRNA-R or IMK3-Sa-sgRNA-F and IMK3-Sa-sgRNA-R, respectively) were annealed and ligated into *pEN-Sa_Chimera*. The entry clones were confirmed by Sanger sequencing, and the inserts transferred into the modified binary vector *pDe-Sa-CAS9* via Gateway *LR* recombination (Thermo Fisher Scientific). Sanger sequencing was performed with primer SS42 (Schiml et al., 2016).

Tubulin marker constructs were previously reported (Schmidt-Marcec et al., 2023). To express *proXVE:IMK2^SP^-GFP and proXVE:GFP-IMK2^SP^*, cDNA corresponding to IMK2 amino acids 1–35 was amplified with primers IMK2-GW-F and IMK2A_B2_R, or IMK2A_B2r_F and IMK2A_B3_R, respectively. Fragments were recombined into Gateway vector *pH7m34* as previously described (Schmidt & Smertenko, 2019). ProCaMV35S:AtFH8-GFP construct was described previously (Zhang et al., 2021).

### Genetics and root analysis

All *A. thaliana* mutant and transgenic lines are listed in **Table S3.** T-DNA lines were provided by ABRC. Two SALK homozygous lines for *IMK2* were isolated: SALK_029864C (*imk2-1*), with an insertion site in the second exon, was genotyped using gene-specific primers imk2F2 and imk2R2 along with left border specific primer Lba1 (**Figure S2A-C**); SALK_081988C (*imk2-2*) with the insertion site in the 5’UTR, was genotyped using gene-specific primers imk2R1 and imk2F1 in combination with primer Lba1. For *IMK3,* the homozygous line SALK_024031C (*imk3-1*), carrying an insertion in the third exon, was genotyped using gene-specific primers imk3F1 and imk3R with Lba1 (**Figure S7A,B**).

To generate double-knockout alleles, *IMK2* was mutagenized in the *imk3-1* background using CRISPR/Cas9 (Schiml et al., 2016) to generate the *imk2-3imk3-1* allele, and *IMK3* was mutagenized in the imk2-2 background using CRISPR/Cas9 to generate the *imk2-2imk3-1* allele. *imk2-3imk3-1* and *Imk2-2imk3-2* T2-generation mutants were validated via Sanger sequencing. T3-generation lines were additionally screened for the lack of *Cas9* gene via PCR. We selected likes that were homozygous for both mutations while lacking the Cas9 vector backbone for phenotypic analysis. *A. thaliana* plants were transformed via floral dip method using *Agrobacterium tumefaciens* strain GV3101 carrying the corresponding construct (Clough & Bent, 1998).

Root growth rate and apical meristem size were measured using seedings grown on vertical agar plates. Seeds were surface-sterilized, sown on half-strength MS (Murashige & Skoog Basal Salt Mixture; Phytotechnology Laboratories, #M524) medium containing 0.7% (w/v) agar, vernalized at 4°C for 48 hours, and then incubated vertically at 22°C with 16-hours light at 100 to 150 mE/m^2^. The root apical meristems were measured in five-day-old seedlings. The root apical meristem was defined from the quiescent centre to the region where cortical cell length exceeds width. For the root growth measurements, three-day-old seedlings were transplanted onto square plates with half-strength MS-medium containing 0.7% (w/v) agar, and the position of the root tips were marked. The seedlings were incubated vertically for a further six days, and the root tips were marked again. The root growth rate was calculated as a length between the two marked dots.

### Chemical genetics

The stock solutions of all drugs were made in DMSO. Arabidopsis roots were grown on vertical agar plates for 5 days (root length 5 to 10 mm). The seedlings were then transferred to 35 mm microscopy petri dishes containing half-strength MS liquid medium supplemented with endosidin 2 at a concentration of 40 µM, endosidin 7 at a concentration of 40 µM, propyzamide at a concentration of 20 µM, brefeldin A at a concentration of 10 µM, concanamycin A at a concentration of 1 µM, caffeine a concentration of 100 µM, or latrunculin B at a concentration of 1 µM for 3 hours. Taxol was added at a concentration of 20 µM for 30 minutes. Roots were then mounted under an agar slab of half strength MS medium with 0.7% agar and imaged immediately. At least five seedlings were imaged per treatment.

BY-2 cell cultures were treated by adding the inhibitor to the growth medium. Propyzamide was added at a concentration of 5 µM, taxol was added at a concentration of 10 µM, and endosidin 7 was added at a concentration of 20 µM. Mock treatments consisted of DMSO at the same final concentration used in drug-supplemented media.

### BY-2 tissue culture and transformation

*N. tabacum* BY-2 cells were maintained according to published procedure (Nagata & Kumagai, 1999, Nagata *et al*., 1981). Generation of stable transgenic lines using *Agrobacterium tumefaciens*-mediated transformation was performed as described previously (Zhang et al., 2021). To produce a cell line expressing *proCaMV35S:IMK2-GFP* and *proCaMV35S:mCherry-TuB6* (Abe & Hashimoto, 2005), *proCaMV35S:mCherry-TuB6* was into BY-2 cell line already expressing *proCaMV35S:IMK2-GFP*. Cell lines expressing both markers were selected under a fluorescence dissecting microscope based on green and red fluorescence intensity. All transgenic BY-2 lines used in this study are listed in **Table S4**.

Cell-cycle synchronization was performed following a published procedure (Zhang et al., 2021). Specifically, a 250 ml flask with 60mL of fresh BY-2 growth medium containing 3 µM aphidicolin was inoculated with 6 mL of seven-day-old BY-2 liquid cell culture. Following 24 hours of incubation under standard growth condition, cells were washed in a sintered glass funnel with a total of 1.5 L 30% (w/v) sucrose solution to remove aphidicolin. After incubating in 60 mL of fresh medium for 4-6 hours, the cells were treated with propyzamide (final concentration 5 µM). Accumulation of metaphase cells was monitored hourly (5-7 hours in total) by staining cell aliquots with DAPI solution as follows: 0.1 mL of BY-2 cells were mixed with 0.5 mL of staining solution (3.7% w/v paraformaldehyde; 50mM PIPES, pH 6.8; 0.4% v/v Triton X-100; 150 nM DAPI) and incubated for 5 minutes at room temperature (Smertenko *et al*., 2016). Specimens were visualized under a Leica DMI4000 epifluorescence microscope equipped with a 40x, NA1.25 oil objective.

Once the metaphase index plateaued, propyzamide was washed out as above and cells were resuspended in 60 mL of fresh medium. Telophase cell accumulation was monitored every 40 minutes using DAPI staining as described. When telophase cell frequency reached 40%, cells were collected by filtering through 50 mesh cloth and flash frozen in liquid nitrogen.

### Bimolecular fluorescence complementation

Full-length *IMK2* and *IMK3* were introduced into Gateway binary vectors as follows. To fuse the gene N-terminally to the N-terminal or C-terminal portion of enhanced yellow fluorescent protein (EYFP), pSITE-nEYFP-N1 and pSITE-cEYFP-N1 were used, respectively, for *IMK2* or *IMK3*. To fuse the gene C-terminally to the C-terminal or N-terminal portion of EYFP, pSITE-cEYFP-C1 (for *IMK3*) and pSITE-nEYFP-C1 (for *IMK2*) were used, respectively (Martin *et al*., 2009). AtFH1 and AtFH8 constructs were described previously (Zhang et al., 2021). Constructs were transformed into *Agrobacterium* strain GV3101. Cultures were grown overnight in YEB medium at 30°C with shaking at 200 rpm shaking until OD_600_ reached 0.5–1.0. Cells were collected by centrifugation for 5 min at 3000 g and washed two times with infiltration medium (10 mM MES, pH 5.6; 10 mM MgCl2; 200 μM acetosyringone). Bacteria were resuspended in 1.0 ml of infiltration medium and incubated at room temperature for 2–5 h. Final OD_600_ value of bacteria solution for infiltration was adjusted to 0.6 with the infiltration medium. Cultures were mixed with cells harbouring p19 plasmid in a ratio of construct1:construct2:p19 = 1:1:0.5 and infiltrated into the abaxial side of *N. benthamiana* leaves. Images were acquired two days after the infiltration as described below.

### Live cell imaging

All IMK2-GFP imaging was performed with *imk2-2;proIMK2:IMK2-GFP#3* line (**Figure S2I**) and IMK3-GFP imaging was performed with *imk3-1;proIMK3:IMK3-GFP#1* line (**Figure S7G**). Roots were mounted under a slab of solid half strength MS medium with 0.7% (w/v) agar on a round 3-cm diameter microscopy dish. Embryos were extracted from developing seeds under a dissecting microscope and mounted in embryo cultivation medium in a round 3-cm diameter microscopy dish under a round coverslip (Lonien & Schwender, 2009). Other plant organs were mounted on a glass microscopy slide in water under a size one coverslip. For cell membrane staining, the mounting medium was supplemented with a freshly prepared 1 μM FM4-64 solution from a 5 mM stock in DMSO. For cell wall staining, the mounting medium was supplemented with a 15 μM propidium iodide solution freshly prepared from a 15 mM stock solution in water. Images were collected using a Leica SP8 confocal laser scanning microscope equipped with a 40x NA1.3 oil immersion objective. GFP was excited at 488 nm, and FM4-64 was excited at 561 nm. All time-lapse images were collected in a single optical plane with a pinhole size of 1.5 Airy units. The 3D reconstructions of the root apical meristem in **Figs. 1H** and **3E** were generated using a Zeiss 980 Airyscan 2 microscope equipped with a Plan-Apochromat 63x, NA1.40 Oil immersion objective DIC f/ELYRA. GFP was excited with a 488 nm laser at 0.5% power, and the emitted light was collected using a 492-546 nm filter.

EosFP photoconversion experiments were performed on either the Leica SP8 or Zeiss 980 Airyscan 2 systems using a 1.5-s pulse of a 405-nm laser set at 2% power. The objective was a 40x NA1.3 oil immersion for the Leica SP8 system or a 63x, NA1.40 Oil immersion objective for the Zeiss 980 system. The photoconverted signal was imaged with excitation at 561 nm and an emission window of 570–640 nm on the Leica SP8 or a 542–636 nm emission filter on the Zeiss 980.

FRAP experiments were performed using the Leica SP8 system. The photobleaching step was performed with a 2–2.6-second pulse of the 488 nm line from the argon laser at 20% power. Signal recovery was imaged using the 488 nm line of the argon laser at 0.5% power and the emission window was 495–590 nm. The pinhole was 1 Airy unit, and the image acquisition rate was of 2–2.6 s per frame.

FLIP experiments were performed using the Leica SP8, a 40x NA1.3 oil immersion objective, and the pinhole was set to 1 Airy unit. The image acquisition rate was 2 s per frame, and bleaching was applied before each frame with 20% power of the 488 nm argon laser in two regions of interest (ROIs) marked as B in **Figure 2A**.

Unless otherwise noted, all images in the manuscript are single optical sections.

### Image analysis and statistics

The xy drift in time-lapse images was corrected using the Fiji StackReg plugin, and kymographs were constructed using the Fiji kymograph plugin. Signal recovery in FRAP and FLIP experiments was quantified in the regions of interest using Fiji (Schindelin *et al*., 2012). The values were exported, and the turnover rates were calculated using single-exponential fit, following published procedure (Chang *et al*., 2005).

The ratio between polymeric and free tubulin was calculated from time-lapse image sequences of cells harboring *proTuB2:EosFP-TuB2*, using frames preceding and following the photoconversion laser pulse. The calculation was based on the equation: (I^ROI3^ − I^ROI1^)/ (I^ROI4^ − I^ROI2^), where I is signal intensity in the four ROIs shown in **Figs. 3L and 10J**.

To calculate the enrichment of the IMK2-GFP or IMK3-GFP signal on the cross wall relative to the lateral wall, the GFP signal in both ROIs (**Figure 4E)** was measured as integrated density using Fiji (Schindelin et al., 2012). Signal enrichment (*E*) was defined as the ratio between the integrated density values at the cross wall (I^C^) and at the lateral wall (I^L^).

The immobile fraction value represents the pool of bleached fluorescent molecules within the FRAP ROI that do not exchange with non-breached molecules during image acquisition. It was calculated as the percentage of the bleached signal that did not recover by the end of the experiment. Signal intensity was corrected by the general photobleaching using a control ROI located in a non-bleached phragmoplast area.

Phragmoplast expansion rate was calculated from 2-3 minutes long time-lapse images taken with 3 seconds frame rate. Signal intensity profiles were generated using the Plot Profile function in Fiji.

Statistical analyses and charts construction were performed using GraphPad Prism 5.0. Statistical comparisons were made using one-way ANOVA or a two-tailed unpaired t-test. For ANOVA, different letters indicate statistically distinct datasets. For *t*-tests, *p* values are provided on each chart for significantly different datasets. Error bars represent standard deviation unless otherwise stated.

### Pull-down assays and proteomics

Cell-cycle synchronized BY-2 cultures were homogenized with a mortar and pestle in liquid nitrogen. Total protein was extracted following a previously described protocol (Avila *et al*., 2015), with the following modifications. One part of the homogenized cell powder was mixed with four parts of Buffer A (100mM Tris-HCl, pH 7.3; 150 mM NaCl; 1 mM EDTA; 10% (w/v) glycerol; 10 µl/ml Leupeptin; 10 µl/ml Pepstatin A; 5 mM AEBSF; 1% Triton X-100 (w/v); 1mM Na_2_MoO_4_; 10mM Na_2_VO_4_; 50mM beta-glycerophosphate; 20mM NaF), and further homogenized with a pestle. After centrifugation for 5 minutes at 20,000 x *g*, 4°C, the supernatant was added to GFP-Trap agarose beads (ChromoTek) pre-washed in Buffer B (100mM Tris-HCl, pH 7.3; 150 mM NaCl; 1 mM EDTA; 10% (v/v) glycerol; 1% (w/v) Triton X-100). The samples were incubated for 2 hours at 4°C with end-over-end rotation. Then the beads were washed in Buffer A twice, and the presence of IMK2-GFP was verified by Western blot using GFP antibodies. Protein composition was analyzed using liquid chromatography–tandem mass spectrometry (LC-MS/MS) followed by peptide analysis at the Southern Alberta Mass Spectrometry Facility, University of Calgary.

### Electron microscopy

Samples for transmission electron microscopy (TEM) were prepared at Washington State University’s Franceschi Microscopy & Imaging Center using microwave assisted tissue fixation with a Pelco BioWave Pro 36500 Laboratory Microwave System (Ted Pella, Inc., Redding, CA, USA). Roots from five-day-old seedlings were fixed in 4% (v/v) glutaraldehyde in a 25 mM cacodylate buffer (pH 7.0) for 5 minutes using microwave conditions set to 150 W and 28°C. Samples were post-fixed in 1% (w/v) osmium tetroxide for 5 minutes under the same microwave settings. After three two-minute rinses in 25 mM cacodylate buffer (pH 7.0), the samples were dehydrated in graded ethanol series from 0% to 50% (v/v) in 10% increments. Following a second rinse with 50% (v/v) ethanol, roots were left in this solution to incubate overnight at -20°C. They were then transferred to 70% (v/v) ethanol (overnight at -20°C), followed by 95% (v/v) ethanol for 24 hours at -20°C. All ethanol solutions were pre-chilled at -20°C. The samples were subsequently washed in absolute ethanol at room temperature, followed by a wash in propylene oxide, and then embedded in Spurr’s low viscosity epoxy resin.

Embedded samples were sectioned using a Leica Ultracut R ultramicrotome equipped with a diamond knife (Leica Microsystems Inc., Deerfield, IL, USA). Sections were stained with 1% (w/v) uranyl acetate and 1% (w/v) lead citrate, and viewed with an FEI Tecnai G2 20 Twin transmission electron microscope (Thermo Fisher Scientific Inc., Waltham, MA, USA). Cell plate morphology was measured on the micrographs with Fiji.

## Acknowledgements

We thank John Sedbrook for sharing *pDe-Sa-CAS9* vector, Chris Staiger for sharing Col-0*;proBRI1:BRI1-GFP* seeds, Michiko Sasabe and Ram Dixit for sharing *map65-1map65-2* seeds, ABRC for providing seeds of T-DNA mutants, Rafal Kacprik for helping with imaging BY-2 cells, Laurent Brechenmacher for help with proteomics analysis, Cecilia Rodriguez-Furlan, Lei Lei, Jamie Van Norman, Georgia Drakakaki, Marcela Rojas-Pierce, Panagiotis Moschou, Keiko Torii, and Ross Sozzani for insightful comments and stimulating discussions.

## Author contributions

Conceptualization and funding acquisition: A.S.; Investigation: T.S., D.F., G.T., A.S.; Methodology: T.S.; Writing – original draft: T.S., A.S.; Writing – reviewing and editing: T.S., D.F., G.T., A.S.

## Data and materials availability

All data are available in the main text, the supplementary materials, or available on request.

---

**Fig. S1.**
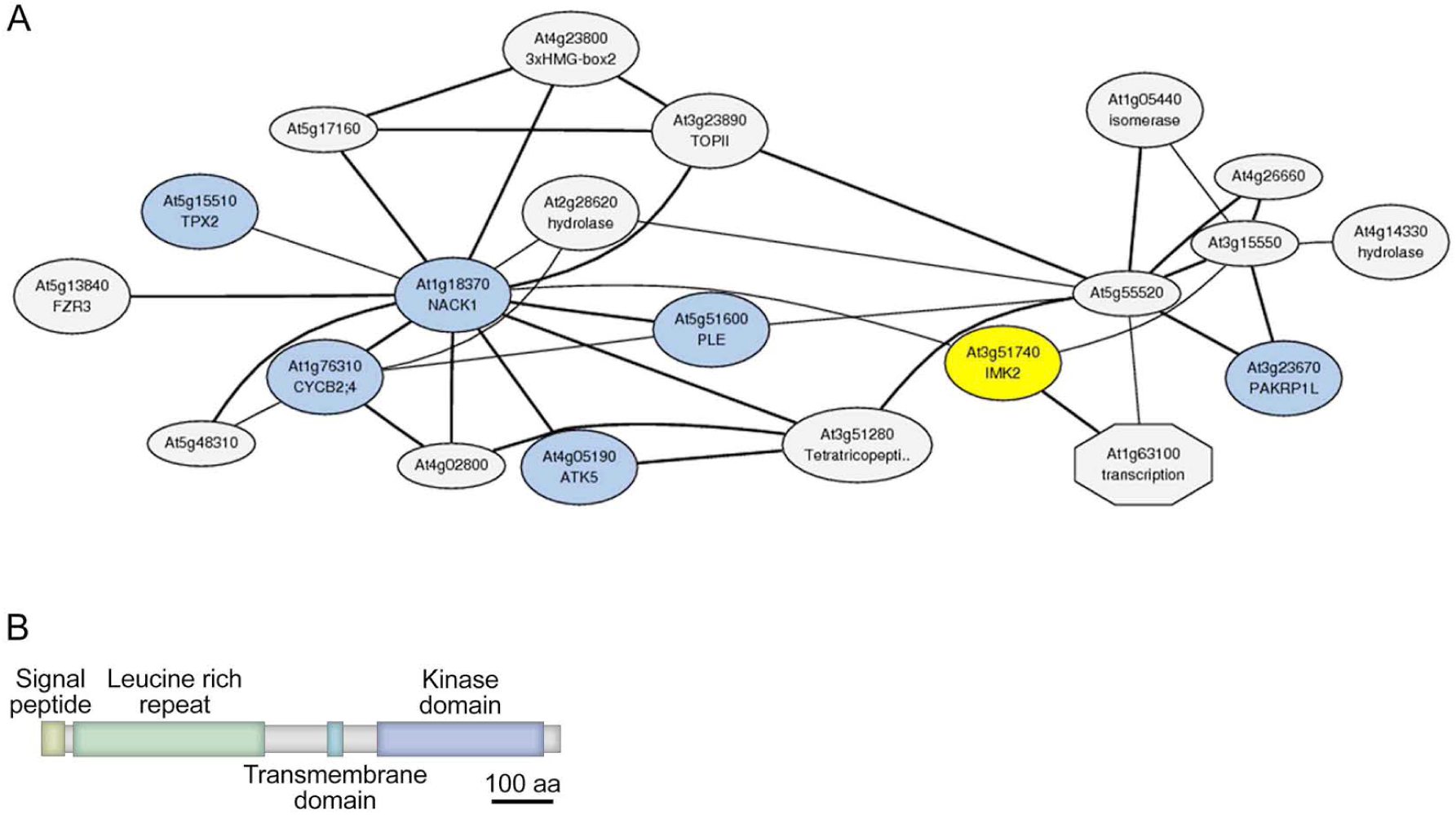
Co-expression analysis of *IMK2*. A, The co-expression network of *IMK2* (highlighted in yellow) includes genes encoding key regulators of cell division (highlighted in blue), such as the kinesins *NACKl (HINKEL),ATKS,* and *PAKRPlL;* the microtubule nucleation factor *TPX2;* the phragmoplast midzone microtubule-bundling protein in *MAP65-3* (PLE); and cyclin B *(CYCB2;4)*. **B,** The predicted domain architecture of IMK2 is characteristic of the leucine-rich repeat receptor-like kinase (LRR­ RLK) superfamily. Scale bar, 100 amino acids.

**Fig. S2.**
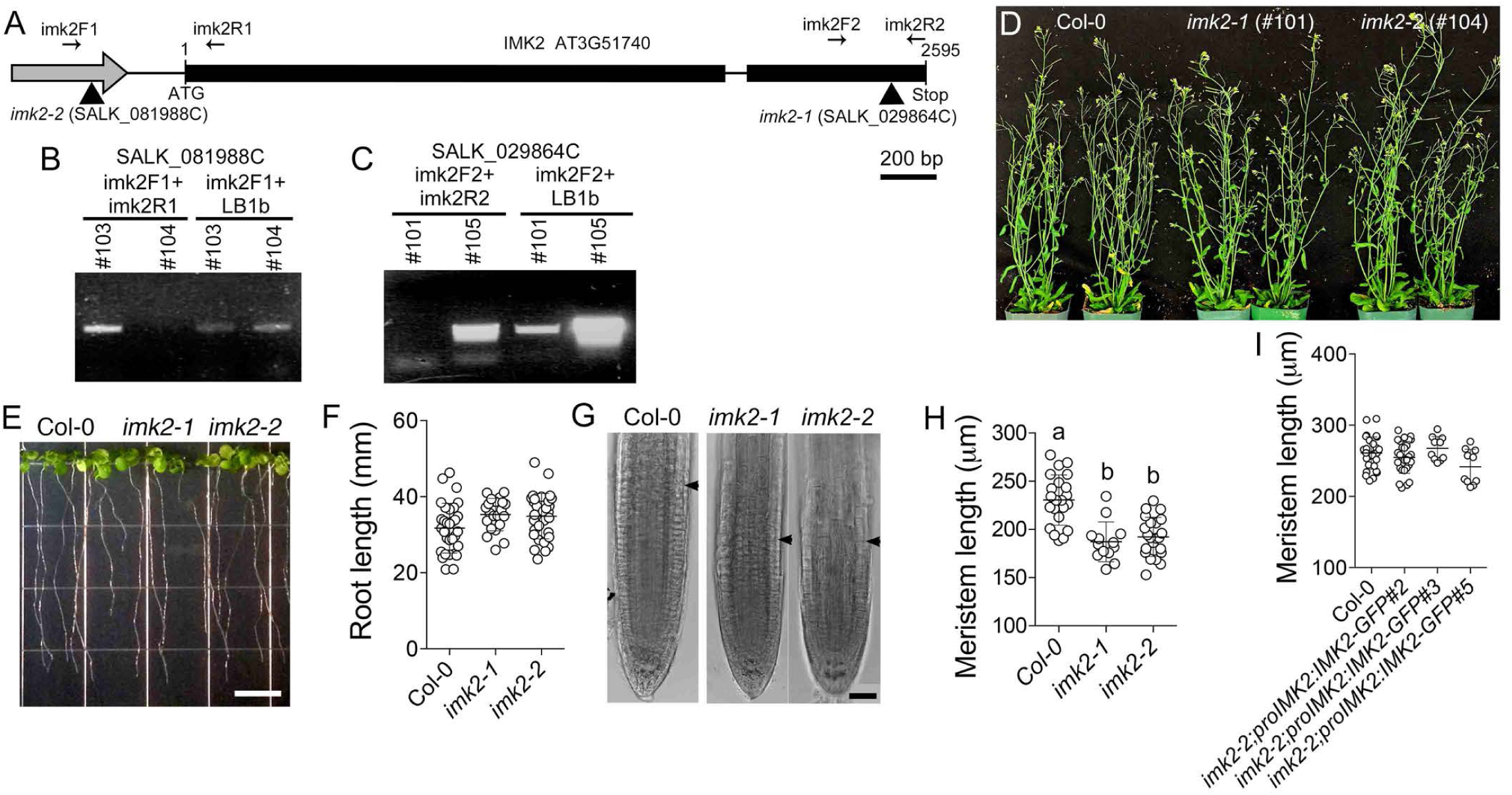
Isolation and characterization of the *IMK2* T-DNA insertion mutants. A, Schematic representation ofT-DNA insertion sites in the *IMK2* gene. The gray arrow indicates the promoter region; thick black lines represent exons, and thin black lines denote introns and non-coding regions. Black arrows indicate the positions of gene-specific primers used for genotyping. The SALK_081988C line *(imk2-2)* contains a T­ ONA insertion 337 bp upstream of the ATG start codon and the SALK_029864C line *(imk2-1)* carries a T-DNA insertion 179 bp upstream of the stop codon. Scale bar, 200 base pairs. **B,C,** Representative images showing PCR-based genotyping of homozygous *imk2* mutants using genomic DNA. In the SALK_081988C line, primers flanking the T-DNA insertion site did not yield a band for plant #104, while a band was detected using the T-DNA left-border primer LBlb and the gene-specific primer imk2Fl, indicating that plant #104 is homozygous **(B).** In contrast, plant#l03 produced bands with both primer pairs, identifying it as heterozygous. Similarly, in the SALK_029864C line, plant#lOl lacked amplification with the T-DNA insertion flanking primers but produced a band using the LBlb and imk2F2 primers, confirming its homozygosity **(C).** Plant #105 showed bands with both primer combinations and was classified as heterozygous. Seeds from plants #101 and #104 were designated *imk2-1* and *imk2-2,* respectively. **D,** Representative images of Col-0, *imk2-1,* and *imk2-2* plants at the flowering stage. **E,** Representative image of roots from nine-day-old Col-0, *imk2-1,* and *imk2-2* seedlings grown vertically. Scale bar, 1 cm. **F,** Root length of Col-0, *imk2-1,* and *imk2-2* seedlings shown in **E.** G, Representative images of root tips in five-day-old Col-0, *imk2-1,* and *imk2-2* seedlings. Arrowheads indicate the distal end of the root apical meristem. Scale bar, 40 µm. **H,** Quantification of meristem length in primary roots of Col-0, *imk2-1,* and *imk2-2* seedlings. Datasets labelled with distinct letters are significantly different (p<0.05) according to one-way ANOVA. I, Apical root meristem length in five-day-old seedlings of Col-0 and three independent *imk2-2* transgenic lines expressing *proIMK2:IMK2-GFP*.

**Fig. S3.**
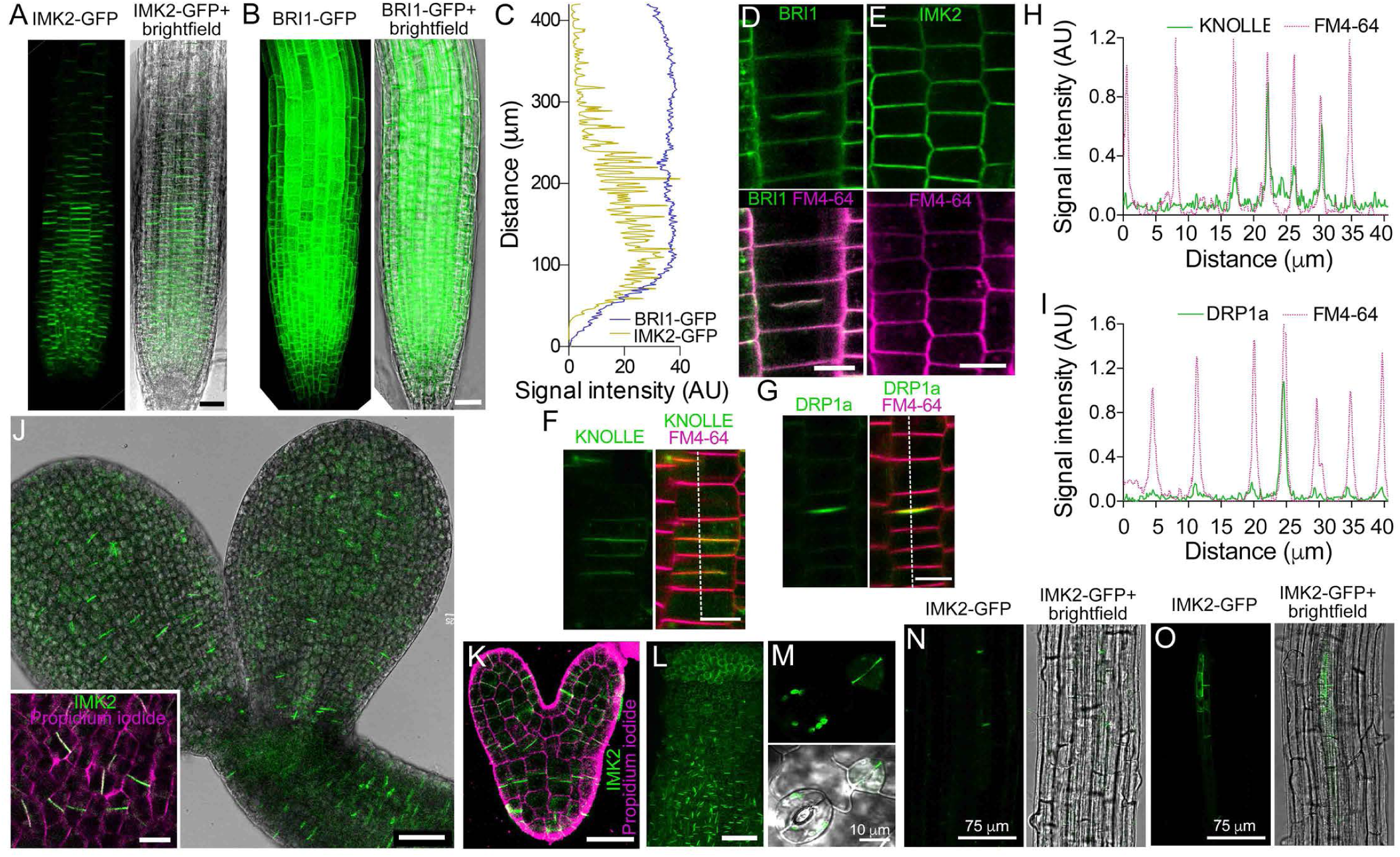
Characterization of IMK2 localization in *A. thaliana*. A, Maximum projection image showing IMK2-GFP localization to the root apical meristem in the *imk2-2;pro/MK2:IMK2-GFP#3* line. Scale bar, 30 µm. This line was used in all IMK2-GFP imaging experiments. **B,** Maximum projection image showing BRil-GFP localization to all cells in the root apical meristem and elongation zone in the *proBR/1:BR/1-GFPline.* Scale bar, 30 µm. **C,** Fluorescence intensity profiles of BRil-GFP and IMK2-GFP measured along the roots shown in panels **A** and **B.** Position 0 on the X-axis corresponds to the tip of the columella cells. **D,** BRil-GFP localizes to the cell plate, lateral walls, and cross walls. Scale bar, 10 µm. **E,** IMK2-GFP localizes to lateral and cross walls in root apical meristem cells of lines expressing *IMK2-GFP* under the control of the constitutive CaMV3SS promoter. Scale bar, 20 µm. **F,** KNOLLE-YFP localization in root apical meristem cells. Scale bar, 10 µm. **G,** DRPla-GFP localization in root apical meristem cells. Scale bar 15 µm. **H,** Normalized fluorescence signals of KNOLLE-YFP and FM4-64 measured along the line shown in panel **F.** I, Normalized fluorescence signals of DRPla-GFP and FM4-64 measured along the line shown in panel **G.** J, Maximum projection image showing IMK2-GFP localization in a mature embryo. Scale bar, 40 µm. Inset shows GFP signal and cell wall staining in cotyledon pavement cells. Scale bar, 10 µm. **K,L,** Maximum projection images showing IMK2-GFP localization in a heart-stage embryo **(K)** and in the epidermis of a developing style **(L).** Scale bars, 25 µm and SO µm, respectively. **M,** IMK2-GFP localizes to the cell wall between recently divided stomata) cells in a developing leaf. Scale bar 10 µm **(M).** N,O, Maximum projection images showing IMK2-GFP localization in the root differentiation zone **(N)** and in lateral root primordia **(0).** Scale bars, 75 µm.

**Fig. S4.**
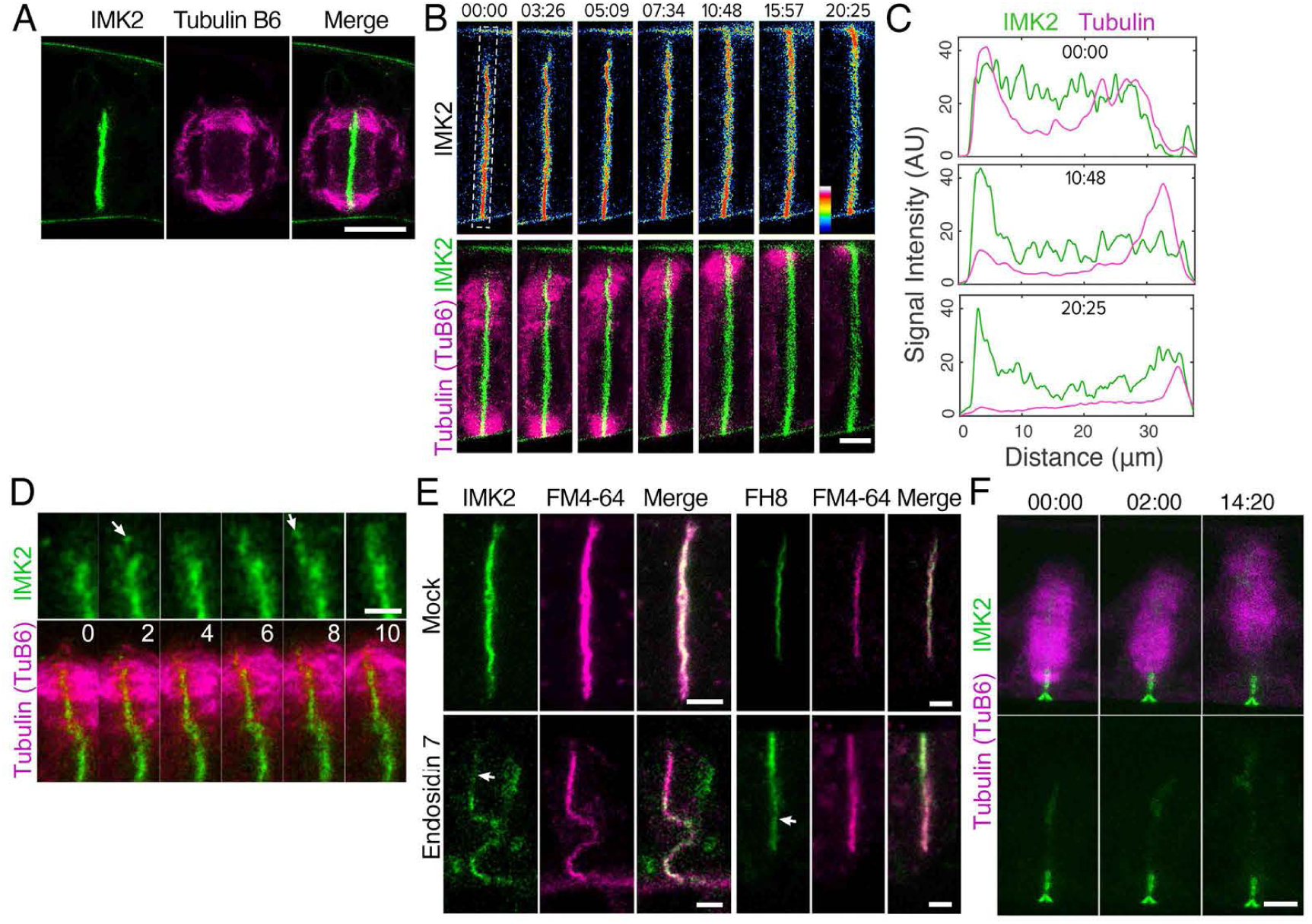
Localization of *A. thaliana* IMK2 in tobacco BY-2 tissue culture cells. A, IMK2-GFP localizes to the phragmoplast midzone. Microtubules are labelled with mCherry-Tu86. Scale bar, 10 µm. **B,** Time-lapse sequence showing the accumulation of IMK2-GFP at the site of cell plate attachment to the parental cell wall. Numbers above images indicate relative time in minutes:seconds. Scale bar, 5 µm. **C,** Fluorescence intensity of IMK2-GFP and mCherry-Tu86 signal in the region highlighted in frame 00:00 at time points 00:00, 10:48, and 20:25. IMK2-GFP signal remains at the cell plate attachment site after the phragmoplast microtubules disappear, as seen in frame 20:25. **D,** Vesicles containing IMK2-GFP (arrows) are delivered to the leading edge of the phragmoplast and subsequently incorporated into the cell plate. Numbers above images indicate relative time in minutes:seconds. Scale bar, 2 µm. **E,** Treatment with endosidin 7 prevents IMK2-GFP accumulation at the phragmoplast midzone, even though a cell plate forms, as revealed by FM4-64 staining. Localization of *A. thaliana* formin FH8 at the phragmoplast midzone is unaffected by endosidin 7. The newly formed sections of the cell plate are indicated by arrows. Scale bar, 5 µm. **F,** Time-lapse frames showing phragmoplast expansion in a cell treated with endosidin 7. IMK2-GFP is delivered to the forming cell plate but dissociates behind the phragmoplast. In contrast, under control conditions (panel **B),** IMK2-GFP remains associated with the cell plate within and behind the lagging zone. Numbers above images indicate relative time in minutes:seconds. Scale bar, 2 µm. Scale bar, 5 µm.

**Fig. S5.**
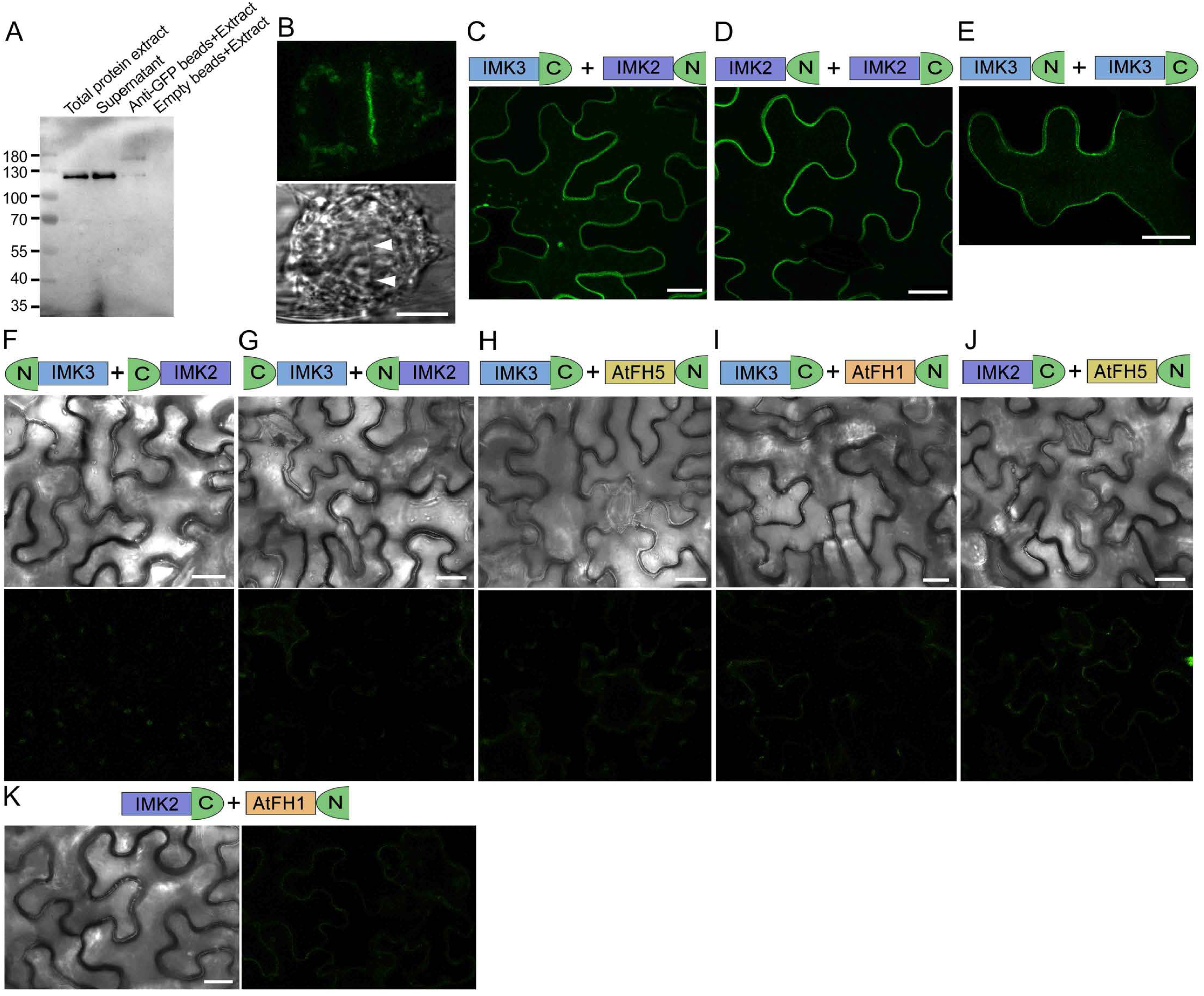
IMK2 interacts with IMK3. A, Western blot analysis with GFP antibodies of samples containing AtIMK2-GFP immunoprecipitations from telophase-synchronized BY-2 cell extracts. Numbers indicate size of molecular markers (kD). **B,** *A. thaliana* IMK3-GFP localizes to the cell plate in BY-2 cells. Scale bar, 10 µm. Arrowheads indicate the position of the cell plate. **C,** Reconstitution ofYFP fluorescence on the plasma membrane in cells co-expressing (-terminal fusion of IMK3 to the C-terminal fragment ofYFP and C-terminal fusion of IMK2 to the N-terminal fragment ofYFP. Image shows a single optical section taken through the central region of the cells. Scale bar, 25 µm. **D,E,** YFP fluorescence reconstitution in cells transiently expressing the indicated protein combinations. Reconstitution of fluorescence confirms that IMK2 and IMK3 interact with each other and with themselves. Scale bars, 25 µm. **F-K,** Negative controls for the BiFC experiments demonstrate that IMK2 or IMK3 do not interact with the integral membrane formins FHl or FH5, nor with each other when the YFP fragments are fused to the N-terminus. In this configuration, the N-terminal YFP fragment masks the signal peptide, preventing proper plasma membrane localization. Protein combinations are indicated above each panel. Scale bars, 25 µm.

**Fig. S6.**
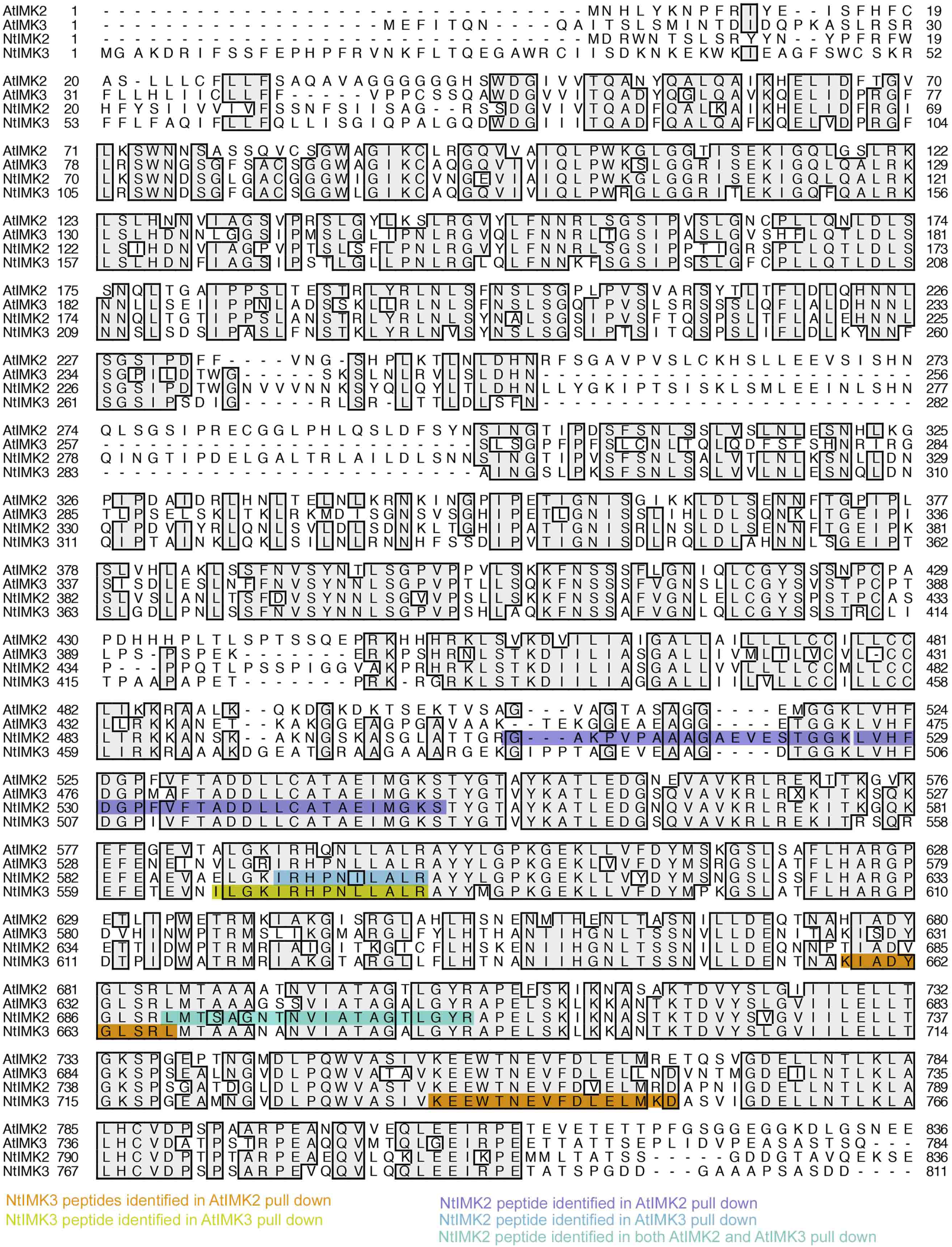
A map of tobacco NtIMK2 and NtIMK3 peptides identified in protein complexes immunoprecipitated with *A. thaliana* AtIMK2 or AtIMK3 from telophase-synchronized extracts of BY-2 cells.

**Fig. S7.**
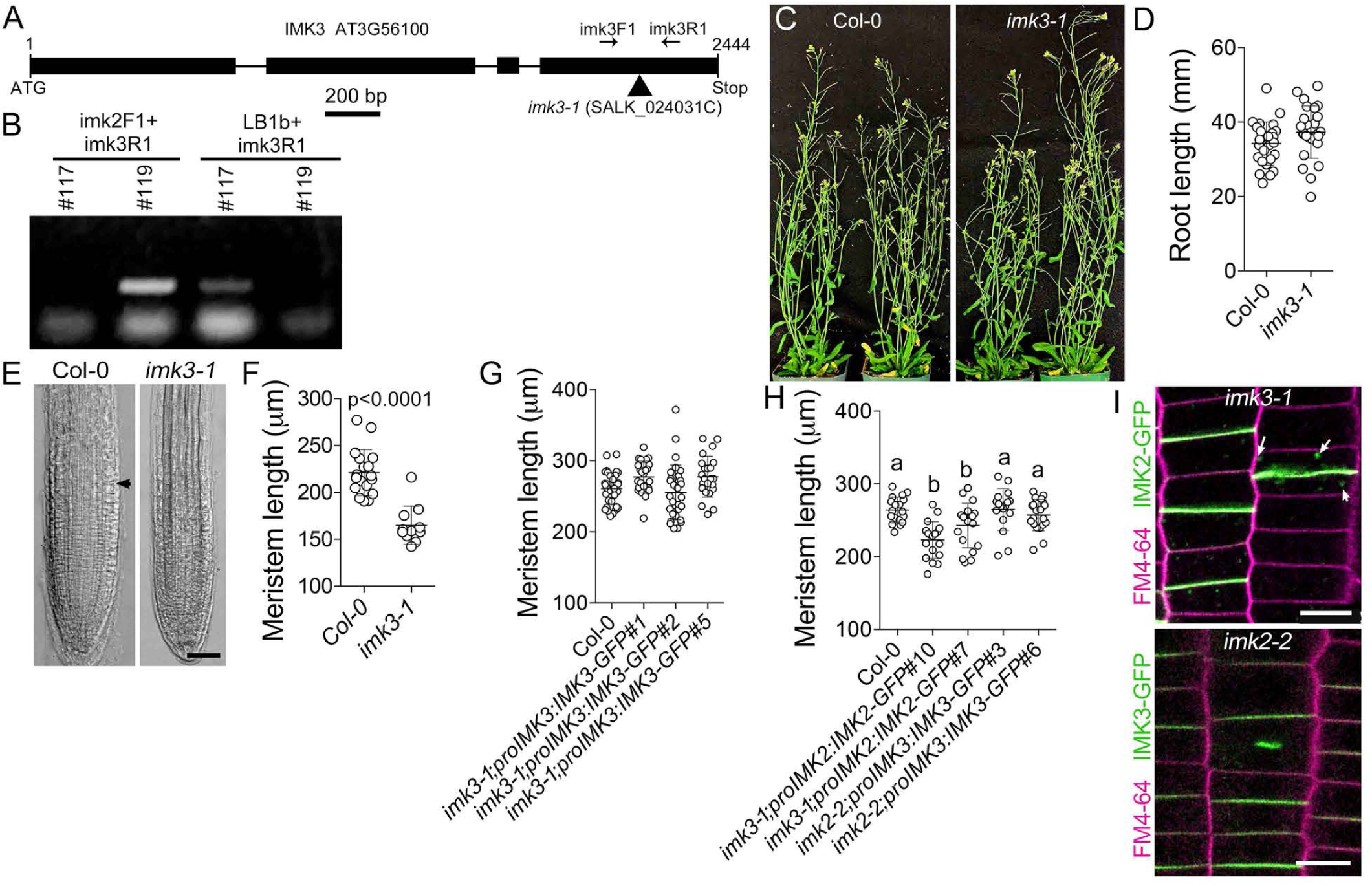
Characterization of the *IMK3* t-DNA insertion mutant. A, Diagram showing the T-DNA insertion site in the *IMK3* gene. The SALK_024031C line contains a T-DNA insertion located 298 bp upstream of the stop codon. Arrows indicate the positions of gene-specific primers used for genotyping. **B,** Identification of the homozygous plant by PCR using genomic DNA. Primers flanking the T-DNA insertion failed to amplify a product in plant #117, whereas amplification with the T-DNA left-border primer (LBlb) and the gene­ specific primer imk3Rl yielded a product, indicating a homologous genotype. In contrast, PCR with DNA isolated from plant #119 produced a band only with the gene-specific primer pair, indicating heterozygosity. Seeds from plant #117 were designated *imk3-1*. **C,** Representative images of Col-0 and *imk3-1* plants during the flowering stage. **D,** Root length of nine-day-old seedlings of Col-0 and *imk3-1* grown vertically. **E,** Representative images of root apical meristem in five-day-old Col-0 and *imk3-1* seedlings. Arrowheads mark the distal end of the root apical meristem. Scale bar, 50 µm. **F,** Quantification of apical root meristem length in five-day-old Col-0 and *imk3-1* seedlings. p-values calculated using an unpaired *t-test* (n<10). **G,** Quantification of apical root meristem length in five-day-old *imk3-1* seedlings transformed with *pro/MK3:IMK3-GFP*. **H,** Quantification of apical meristem length in five-day-old *imk3-1* complemented with *pro/MK2:IMK2-GFP* and in *imk2-2* complemented with *pro/MK3:IMK3-GFP.* Datasets labelled with distinct letters are significantly different (p<0.05) according to one-way ANOVA. I, Localization of IMK2-GFP in the *imk3-l;pro/MK2:IMK2-GFP#l0* line and IMK3-GFP in the *imk2-2;pro/MK3:IMK3-GFP#6* line. Arrows point to vesicles containing IMK2-GFP. Scale bars, 10 µm.

**Fig. S8.**
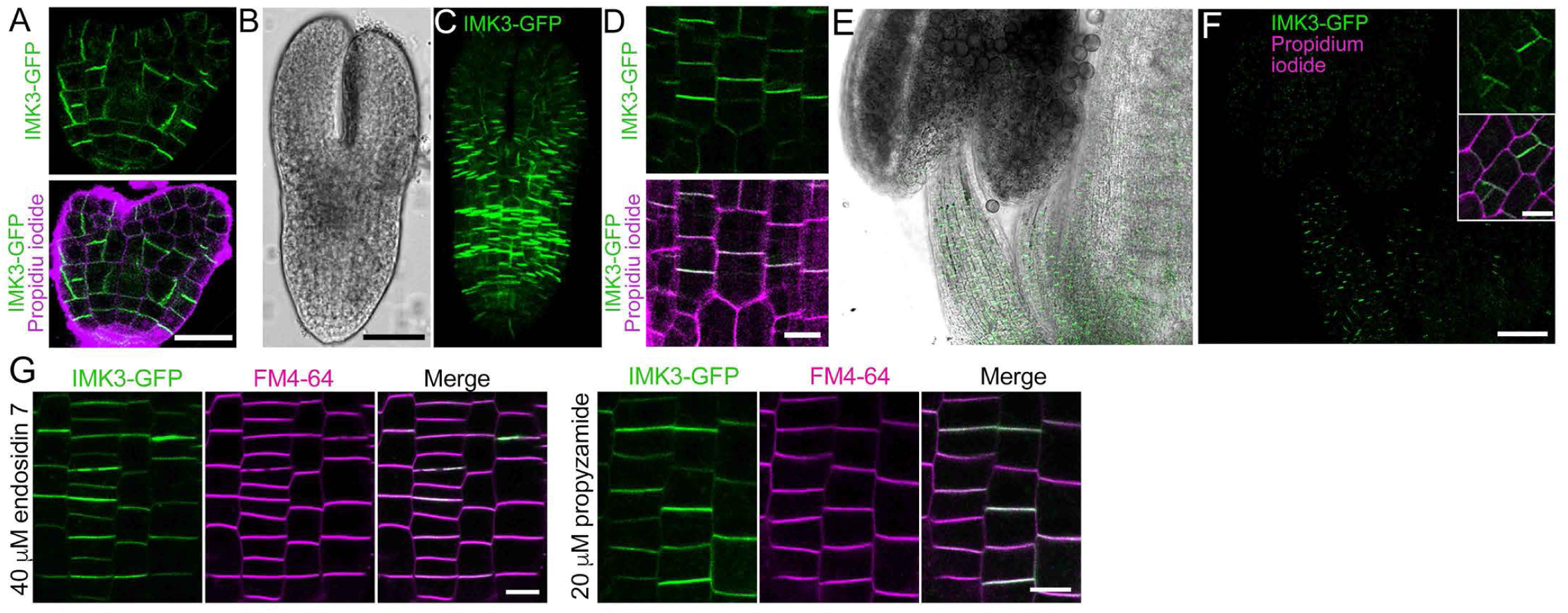
Analysis of IMK3-GFP localization. A, IMK3-GFP localization in a heart-stage embryo of the *imk3-1;pro/MK3:IMK3-GFP#1* line. Scale bar 20 µm. All subsequent images were acquired using this line. **B,C,** Bright field **(B)** and maximum projection IMK3-GFP **(C)** images of a torpedo-stage embryo. Scale bar, 40 µm. **D,** Single optical section of cotyledon epidermal cells from the embryo shown in panel **C.** Scale bar, 10 µm. **E, F,** Maximum projection image showing IMK3-GFP localization in the anther and style of a developing flower. Scale bar 50 µm. The inset in panel **F** shows a single optical section of the style epidermis. Scale bar, 10 µm. **G,** IMK3-GFP localization in root apical meristem cells is not affected by a three-hour treatment with 40 µM endosidin 7 or 20 µM propyzamide. Scale bars, 10 µm.

**Fig. S9.**
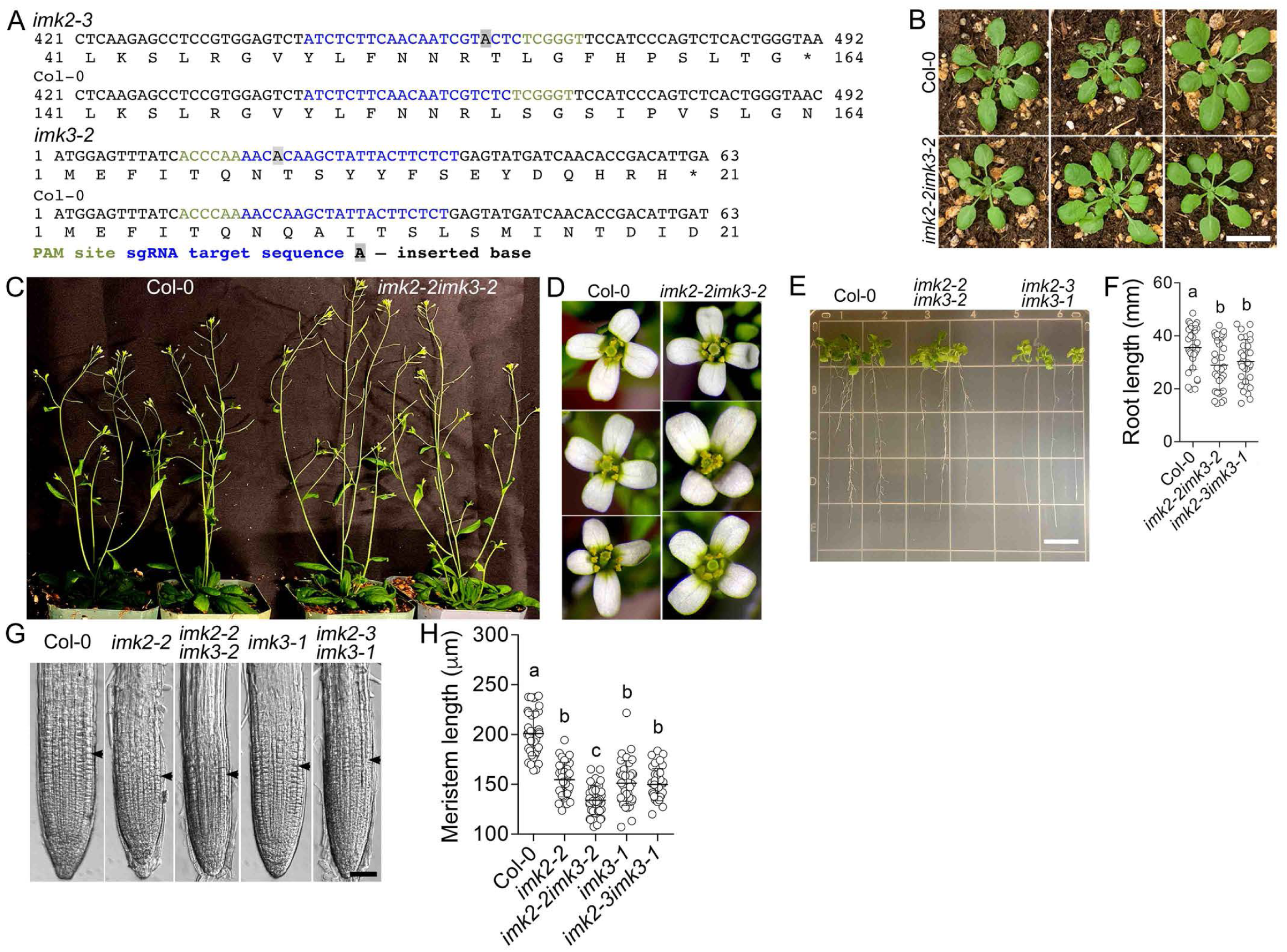
Generation and phenotypic analysis of *IMK2* and *IMK3* double knockout mutants. A, Positions of CRISPR-Cas9-induced insertions in the *IMK2* and *IMK3* genes, and corresponding changes in their reading frames. **B-D,** Representative images of seedlings, mature plants, and flowers of Col-0 and the *imk2-2imk3-2* allele. Scale bar, 1 cm. **E,** Representative images of Col-0, *imk2-2imk3-2,* and *imk2-3imk3-1* seedlings grown on vertical half-strength MS agar plates. Scale bar, 1 cm. **F,** Quantification of Col-0, *imk2-2imk3-2,* and *imk2-3imk3-1* root length. Datasets labelled with distinct letters are significantly different (p<0.05) according to one-way ANOVA (30 roots grown on five separate plates were measured per each genotype). **G,** Representative images of primary root tips in five-day-old seedlings of Col-0, single, and double mutants. Arrowheads mark the distal end of the meristem. Scale bar, 50 µm. **H,** Quantification of meristem size in five-day-old seedlings of Col-0, single, and double mutants. Datasets labelled with distinct letters are significantly different (p<0.05) according to one-way ANOVA (over 32 roots grown on three separate plates were measured).

**Fig. S10.**
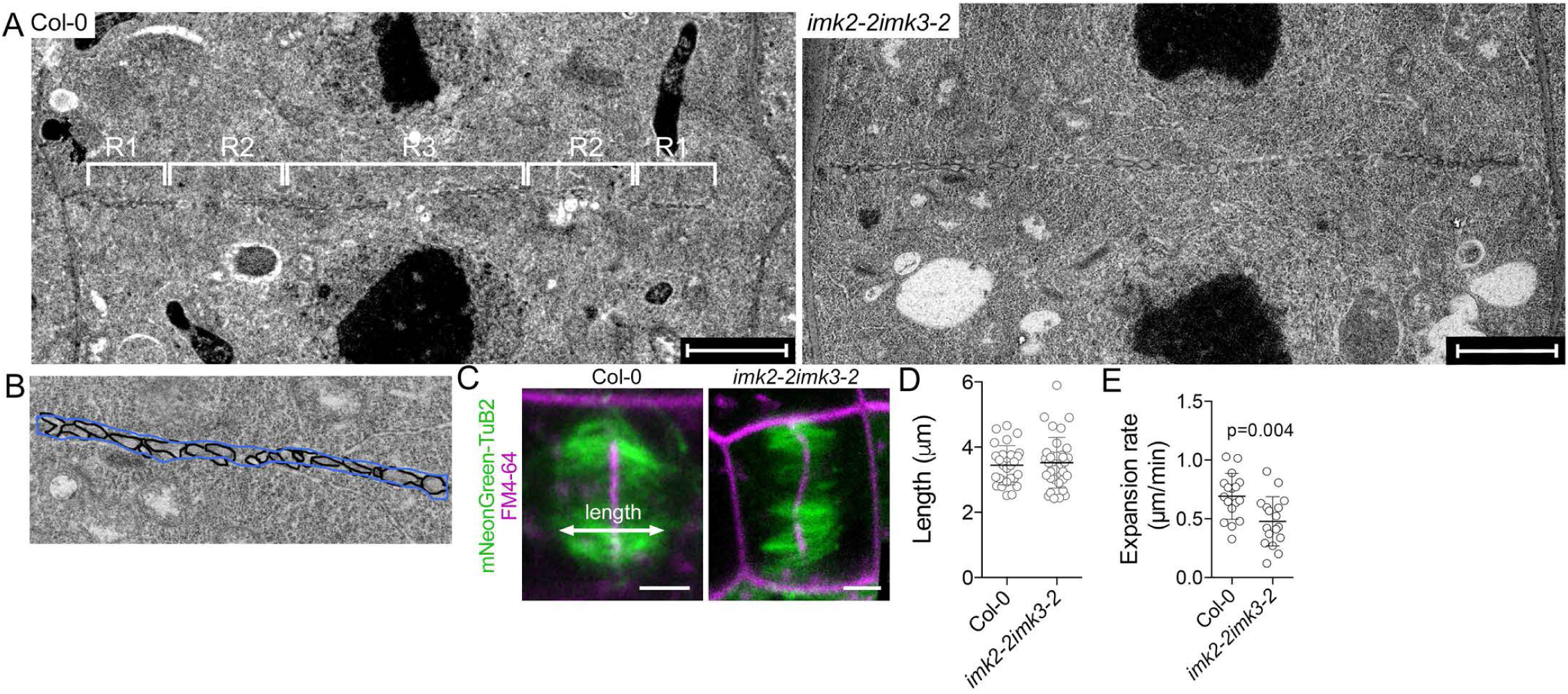
Analysis of cell division in *imk2-2imk3-2*. A, Representative transmission electron micrographs of cell plate in dividing root apical meristem cells of Col-0 and *imk2-2imk3-2* plants used for measuring cell plate compartments. Sections were taken through the central region of daughter cells, showing both daughter nuclei. Regions where the cell plate was measured are marked as Rl, R2, and R3. Scale bars, 2 µm. **B,** Cartoon illustrating how the area occupied by the cell plate compartments was measured. The compartment area was measured as the ROI outlined in blue, and the total cell plate area as the ROI outlined in black The percentage of cell plate area occupied by the compartments was then calculated. **C,** Representative images of phragmoplasts in root apical meristem cells of Col-0 and *imk2-2imk3-2* seedlings. Scale bars, 2.5 µm. **D,** Phragmoplast length in Col-0 and *imk2-2imk3-2,* measured as shown in panel **C.** E, Phragmoplast expansion rate in Col-0 and *imk2-2imk3-2.* p-values calculated using an unpaired *t-test; n=l* 7 (individual phragmoplasts).

**Fig. S11.**
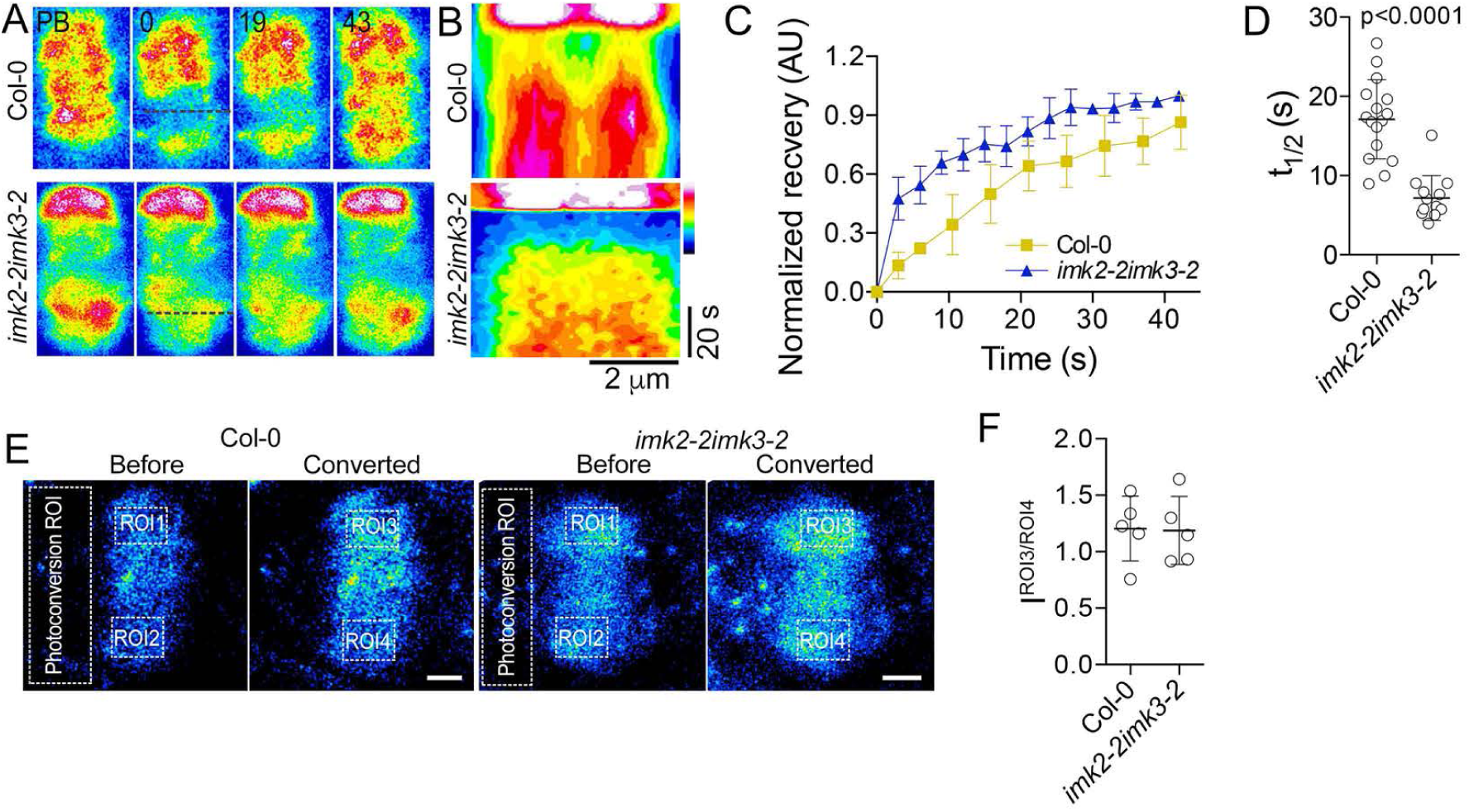
Comparison ofmicrotubule dynamics in Col-0 and *imk2-2imk3-2* phragmoplasts. A, Frames from representative FRAP experiments in root apical meristem cells of Col-0 and *imk2-2imk3-2* expressing mNeonGreen-Tu82. Numbers indicate time after photobleaching (in seconds). The pre-bleach (PB) frame shows the phragmoplast before photobleaching. **B,** Kymographs taken along the black lines in panel **A.** C, Recovery of fluorescent signal in the photobleached ROIs. Error bars denote standard deviation of the mean, *n=3* cells. **D,** Microtubule turnover values (t1;2) in phragmoplasts of Col-0 and *imk2-2imk3-2.* p-values calculated using an unpaired *t-test; n=16* (individual phragmoplasts). **E,** Representative images of EosFP-TuB2 before and after photoconversion with a 405 nm laser pulse. Photoconversion was performed in a cytoplasmic ROI outside the phragmoplast, and the signal was measured in ROis 1 to 4. Scale bars, 2 µm. **F,** Ratio of signal intensities in ROI3 to ROI4 (J^ROI3/ROI4^) following photoconversion of cytoplasmic tubulin in the “Photoconversion ROI” as shown in panel **E.** The following equation was used: J^ROI3/ROI4^=(I^ROI3^-I^ROI1^)/(I^ROI4^-I^ROI2^). The resulting values are close to 1 in both Col-0 and the *imk2-2imk3-2* allele, indicating a similar abundance of photoconverted tubulin in ROI3 and ROI4.

**Table S1.**
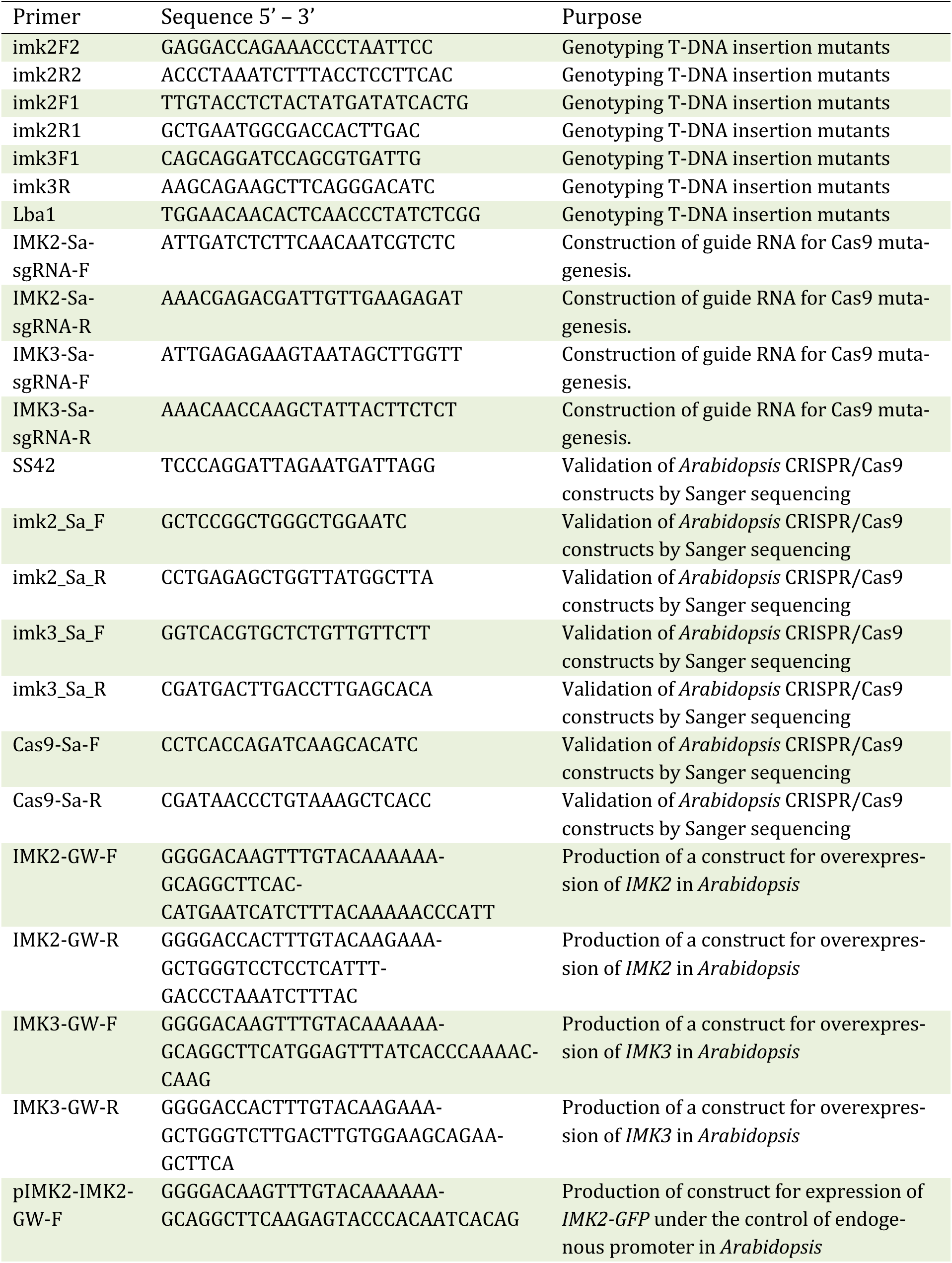
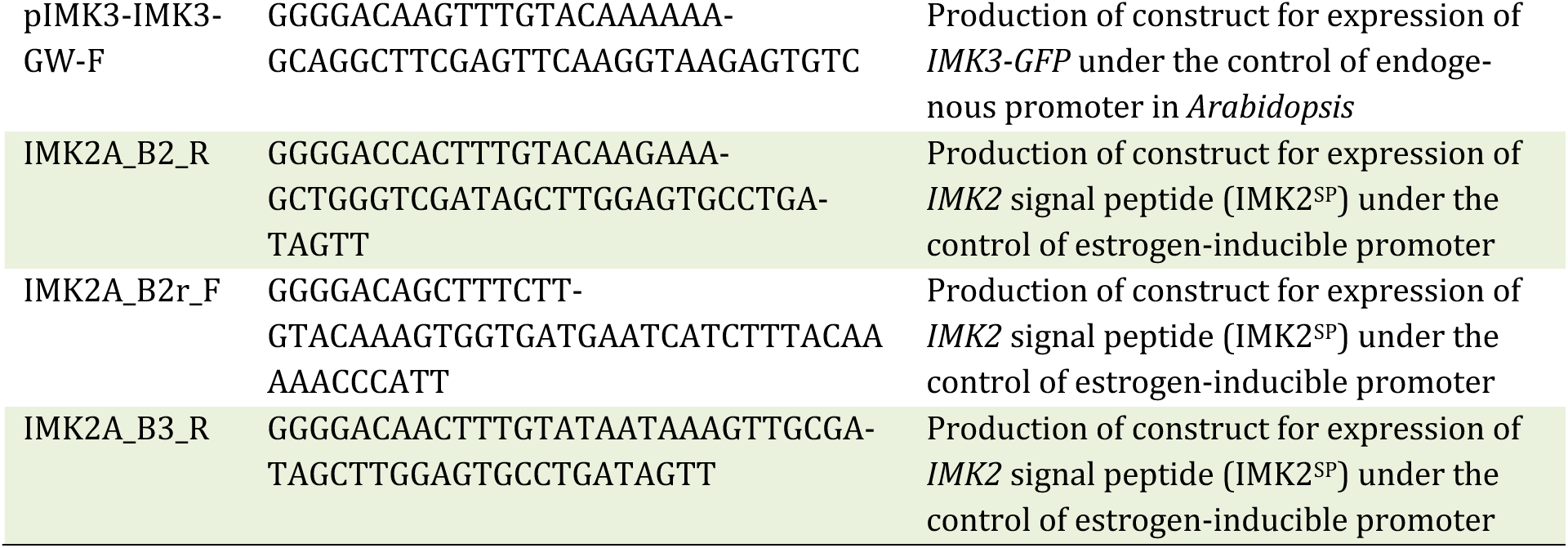
List of primers used in this study.

**Table S2.** List of constructs used in this study.

| Construct | Description | Reference |
| --- | --- | --- |
| *proIMK2:IMK2-GFP* | IMK2 C-terminal fusion to GFP under the control of endogenous promoter | This work |
| *proIMK3:IMK3-GFP* | IMK3 C-terminal fusion to GFP under the control of endogenous promoter | This work |
| *proTuB2:mNeonGreen-TuB2* | β-tubulin 2 N-terminal fusion to mNeonGreen under the control of endogenous promoter | (Schmidt-Marcec et al., 2023) |
| *proTuB2:EosFP-TuB2* | β-tubulin 2 N-terminal fusion to EosFP under the control of endogenous promoter | (Schmidt-Marcec et al., 2023) |
| *proCaMV35S:IMK2-GFP* | IMK2 C-terminal fusion to GFP under the control of cauliflower mosaic 35S (*CaMV35S*) promoter | This work |
| *proCaMV35S:IMK3-GFP* | IMK3 C-terminal fusion to GFP under the control of cauliflower mosaic 35S (*CaMV35S*) promoter | This work |
| *proXVE:IMK2<sup>SP</sup>-GFP* | IMK2 signal peptide with C-terminal GFP under the control of estrogen-inducible promoter | This work |
| *proXVE:GFP-IMK2<sup>SP</sup>* | IMK2 signal peptide of IMK2 with N-terminal GFP under the control of estrogen-inducible promoter | This work |
| *proCaMV35S:mCherry-TuB6* | N-terminal tubulin β6 fusion to mCherry under the control of cauliflower mosaic 35S (*CaMV35S*) promoter | (Abe & Hashimoto, 2005) |
| *IMK2/pSITE-nEYFP-N1* | Full length *IMK2* fused N-terminally to N-terminal portion of enhanced yellow fluorescent protein (EYFP) | This work |
| *IMK2/pSITE-cEYFP-N1* | Full length *IMK2* fused N-terminally to C-terminal portion of EYFP | This work |
| *IMK2/pSITE-nEYFP-C1* | Full length *IMK2* fused C-terminally to N-terminal portion of EYFP | This work |
| *IMK2/pSITE-cEYFP-C1* | Full length *IMK2* fused C-terminally to C-terminal portion of EYFP | This work |
| *IMK3/pSITE-nEYFP-N1* | Full length *IMK3* fused N-terminally to N-terminal portion of EYFP | This work |
| *IMK3/pSITE-cEYFP-N1* | Full length *IMK3* fused N-terminally to C-terminal portion of EYFP | This work |
| *IMK3/pSITE-nEYFP-C1* | Full length *IMK3* fused C-terminally to N-terminal portion of EYFP | This work |
| *IMK3/pSITE-cEYFP-C1* | Full length *IMK3* fused C-terminally to C-terminal portion of EYFP | This work |
| *AtFH5/pSITE-nEYFP-N1* | Full length *AtFH5* fused N-terminally to N-terminal portion of EYFP | (Zhang et al., 2021) |
| *AtFH1/pSITE-nEYFP-N1* | Full length *AtFH1* fused N-terminally to N-terminal portion of EYFP | (Zhang et al., 2021) |

**Table S3.** Arabidopsis lines used in this study.

| Line | Description | Reference |
| --- | --- | --- |
| *imk2-1* | *IMK2* T-DNA knockout allele | This work |
| *imk2-2* | *IMK2* T-DNA knockout allele | This work |
| *imk3-1* | *IMK3* T-DNA knockout allele | This work |
| *imk2-2imk3-2* | *IMK2IMK3* double mutant allele | This work |
| *imk2-3imk3-1* | *IMK2IMK3* double mutant allele | This work |
| *club-2* | Knockout allele of TRAPP<sup>II</sup> subunit TRS130 | (Rybak et al., 2014) |
| *map65-1map65-2* | *MAP65-1* and *MAP65-2* double mutant allele | (Sasabe et al., 2011) |
| Col-0;*proBRI1:BRI1-GFP* | BRI1-GFP under the control of endogenous promoter in Col-0 wild type background | (Geldner et al., 2007) |
| *imk2-2;proIMK2:IMK2-GFP* | *proIMK2:IMK2-GFP* in *imk2-2* background | This work |
| *imk3-1;proIMK3:IMK3-GFP* | *proIMK3:IMK3-GFP* in *imk3-1* background | This work |
| *imk3-1;proIMK2:IMK2-GFP* | *proIMK2:IMK2-GFP* in *imk3-1* background | This work |
| *imk2-2;proIMK3:IMK3-GFP* | *proIMK3:IMK3-GFP* in *imk2-2* background | This work |
| *imk2-3imk3-2;proTuB2:mNeonGreen-TuB2* | *proTuB2:mNeonGreen-TuB2* in *imk2-3imk3-2* background | This work |
| *imk2-3imk3-2;proTuB2:EosFP-TuB2* | *proTuB2:EosFP-TuB2* in *imk2-3imk3-2* background | This work |
| Col-0;*proTuB2:EosFP-TuB2* | *proTuB2:EosFP-TuB2* in Col-0 wild type background | (Schmidt-Marcec et al., 2023) |
| Col-0;*proTuB2:mNeonGreen-TuB2* | *proTuB2:mNeonGreen-TuB2* in Col-0 wild type background | (Schmidt-Marcec et al., 2023) |
| *club-2;proIMK2:IMK2-GFP* | *proIMK2:IMK2-GFP* in *club-2* background | This work |
| *club-2;proIMK3:IMK3-GFP* | *proIMK3:IMK3-GFP* in *club-2* background | This work |
| *map65-1map65-2;proIMK2:IMK2-GFP* | *proIMK2:IMK2-GFP* in *map65-1map65-2* background | This work |
| *map65-1map65-2;proIMK3:IMK3-GFP* | *proIMK3:IMK3-GFP* in *map65-1map65-2* background | This work |
| Col-0;*proCaMV35S:IMK2-GFP* | *proCaMV35S:IMK2-GFP* in Col-0 wild type background | This work |
| Col-0;*proXVE:IMK2<sup>SP</sup>-GFP* | *proXVE:IMK2<sup>SP</sup>-GFP* in Col-0 wild type background | This work |
| Col-0;*proXVE-GFP-IMK2<sup>SP</sup>* | *proXVE-GFP-IMK2<sup>SP</sup>* in Col-0 wild type background | This work |
| Col-0;*proKNOLLE:KNOLLE-YFP* | *KNOLLE-YFP* expressed in Col-0 wild type background under control of the endogenous promoter | (Steiner et al., 2016) |
| Col-0;*proDRP1a:DRP1a-GFP* | *DRP1a-GFP* expressed in Col-0 wild type background under control of the endogenous promoter | (Kang et al., 2003) |

**Table S4.**
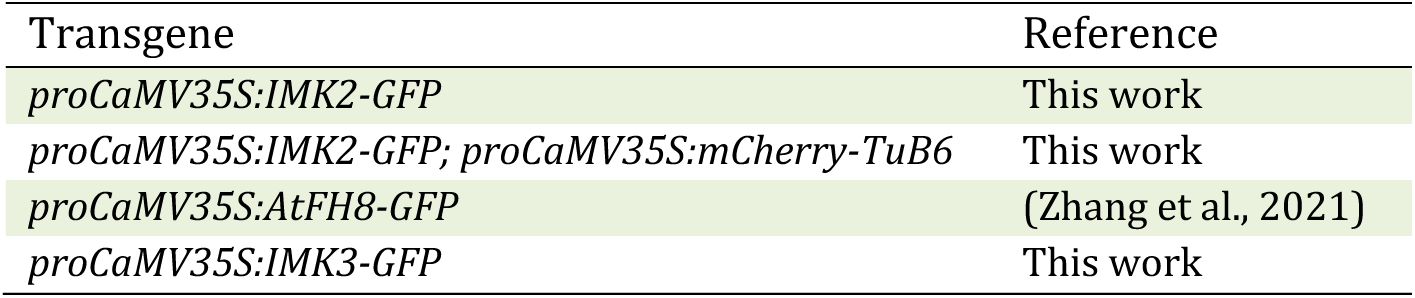
Stable transgenic BY-2 cell lines used in this study.

**Table S5.** Accession number of all genes used in this study.

| Gene name | Accession number |
| --- | --- |
| *IMK2* | At3G51740 |
| *IMK3* | At3G56100 |
| *Formin homology FH1* | AT3G25500 |
| *Formin homology FH5* | AT5G54650 |
| *Formin homology FH8* | AT1G70140 |
| *Beta-tubulin 2 (TuB2)* | AT5G62690 |
| *Beta tubulin 6 (TuB6)* | AT5G12250 |
| *TRS130* | AT5G54440 |
| *DRP1a* | AT5G42080 |
| *KNOLLE* | AT1G08560 |
| *BRI1* | AT4G39400 |
| *MAP65-1* | AT5G55230 |
| *MAP65-2* | AT4G26760 |

